# Bramble: projection of spliced genomic alignments into transcriptomic space for improved transcript quantification

**DOI:** 10.64898/2026.09.09.750464

**Authors:** Zoe Rudnick, Ales Varabyou, Rob Patro, Mihaela Pertea

## Abstract

Accurate transcript abundance estimation is central to many transcriptomic studies. Many current quantification methods rely on reads mapped directly to the transcriptome, but transcriptome alignment can misassign reads from unannotated transcripts to annotated isoforms, leading to biased abundance estimates. We introduce Bramble, a method that projects spliced genomic alignments into transcriptomic coordinates to produce alignments compatible with downstream transcript quantification tools. Across simulated short- and long-read RNA-seq datasets and multiple levels of reference annotation completeness, incorporating Bramble into quantification pipelines consistently improved accuracy and reduced error. These results suggest that genome-derived transcriptomic alignments can improve transcript quantification by preserving compatible alignments to annotated transcripts while filtering alignments likely originating from unannotated transcripts.

## Introduction

Transcript quantification is a central component of RNA-seq analysis, responsible for the estimation of known transcript abundances and depended upon by downstream applications including differential expression analyses and trait association studies. To ensure meaningful downstream results, accurate transcript abundance estimates are crucial. While quantification methods have advanced substantially over the past two decades, current methods still face considerable limitations.

Several strategies exist for quantifying short-read RNA-seq data. One prominent strategy aligns RNA-seq reads to the transcriptome^1–4^ and quantifies abundances using these alignments^5^. Full alignment of every read can be computationally demanding, however, which has prompted the development of lightweight *k*-mer-based mapping techniques^6–9^. When paired with advances like selective alignment^10–11^, lightweight mapping offers significant improvements in runtime while maintaining accuracy.

The adoption of long-read sequencing technologies, including Oxford Nanopore Technologies^12–13^ (ONT) and Pacific Biosciences^14–15^ (PacBio), for RNA-seq applications has motivated the development of alignment^16^ and quantification^17–19^ tools tailored to long-read transcriptomic data. Quantification of long-read data is commonly alignment-based, because, depending upon the protocol and technology, long-read data can display heterogeneous error profiles that often require alignment-level scoring to assess accurately.

A significant limitation faced by current quantification methods is annotation bias, which is caused by a reliance on transcriptome mappings. When reads are aligned or mapped solely to the transcriptome, which comprises sequences described by a reference annotation, reads originating from unannotated transcripts may be misassigned to annotated transcripts, thus skewing abundance estimates^11, 20^. Notably, there is no consensus on a single definitive human transcriptome annotation^21–25^, let alone the annotations of non-model organisms, for which annotation is largely an automated process^26–27^.

Across species, pervasive transcriptional noise caused by low-level, stochastic transcriptional events^28–33^ accounts for almost one-third of cellular RNA molecules^34–35^ and complicates genome annotation by necessitating that annotation methods distinguish between functional and noisy transcripts. As such, it is expected that for any transcriptome, there exist some functional transcripts that have not yet been incorporated and some non-functional transcripts that have been incorrectly incorporated. When annotations are created by automated processes, it is particularly difficult to annotate species-specific genes, leaving many functional transcripts unannotated^26, 36–37^. Furthermore, genes subject to rapid sequence diversification—including viral genes, somatically mutated cancer genes, and adaptive immune-receptor genes—can produce transcripts that differ substantially from the reference, both across individuals and across cells within an individual, making fixed annotations unreliable^38–40^.

We introduce Bramble, a method that projects spliced genomic alignments into transcriptomic space. When RNA-seq reads are aligned to the genome and then projected with Bramble, quantification methods can estimate transcript abundances using genomic alignments, reducing annotation bias caused by transcriptome alignment. With this approach, reads best explained by unannotated loci are not constrained to the space of annotated transcripts. Instead, they can align to the correct loci and be filtered appropriately, preventing skewing of abundance estimates. Bramble also enables splice-informed quantification by incorporating splice junction adherence into assignment and scoring modules, which is not possible with transcriptomic alignments, wherein splice information is not retained. Furthermore, comprehensive reporting of possible transcriptomic alignments is necessary for accurate quantification^41^, but can be prohibitively costly with transcriptome mapping, because searching for secondary alignments requires many successive alignment processes. Bramble solves this problem by determining the set of all compatible transcriptomic alignments simultaneously for every genomic alignment.

At present, there are several tools for short-read data that leverage genome alignment information for transcriptomic analysis. One such tool is STAR^42^, which contains an internal module that projects its own genomic alignments to transcriptomic space, enabling users to obtain genomic and transcriptomic alignments at the same time. However, this method cannot be applied to genomic alignments generated by other aligners. Another is Salmon, which recommends adding decoys derived from the genome sequence to its index to improve quantification accuracy. To the best of our knowledge, no strategy has been developed to date to improve transcriptome-space quantification from long-read data using genome-alignment information. Generally, quantification based on spliced genomic alignments has not yet been systematically assessed in the context of unannotated transcripts and transcriptional noise, which are prevalent in real RNA-seq data but frequently absent in benchmarking studies. Here, we evaluate the performance of Bramble-based quantification pipelines using simulated short- and long-read data containing unannotated transcripts and transcriptional noise, demonstrating consistent improvements over conventional pipelines across read types and levels of reference annotation completeness.

## Results

### Evaluation of performance on short-read data

To benchmark the performance of Bramble on short-read data, we used the simulated short-read dataset and human reference annotation from Varabyou et al. 2021, which includes added transcriptional noise to mimic real RNA-seq data (see “Simulated data”). We first compared transcriptomic alignments produced by HISAT2^43^ + Bramble against alignments produced by Bowtie2, a widely used transcriptome alignment baseline. We ran Bowtie2 with parameters set up to enable near-comprehensive secondary alignment reporting, permitting up to 76 distinct alignments per read; see “Secondary alignments in transcriptome alignment” for details. Command line settings for each tool are described in Supplementary Note 1. We used three metrics to evaluate the alignments: precision, recall, and F1 score. Per-read precision was computed as the number of correct alignments divided by the total number of reported alignments, where alignments are considered correct if they report the true transcript of origin. Per-read recall was set to 1 if the true transcript of origin was reported by at least one alignment and 0 otherwise. We averaged precision and recall scores across reads and samples, binning by the number of isoforms belonging to the gene of origin of each read. For every bin, we calculated the F1 score on the average precision and recall.

Reads from genes with few isoforms had the highest precision, and precision declined as isoform count increased, reflecting the greater number of plausible alignment targets at more complex loci. At every bin, HISAT2 + Bramble achieved higher precision than Bowtie2 (Fig. 1a). Recall was consistently high (>95%) for both methods, with Bowtie2 slightly outperforming HISAT2 + Bramble (Supplementary Fig. 1a). F1 scores were higher for HISAT2 + Bramble (Supplementary Fig. 1b). Our results demonstrate that HISAT2 + Bramble approaches the high recall of Bowtie2 with near-comprehensive secondary alignment reporting, while improving precision at the read level.

**Figure 1.**
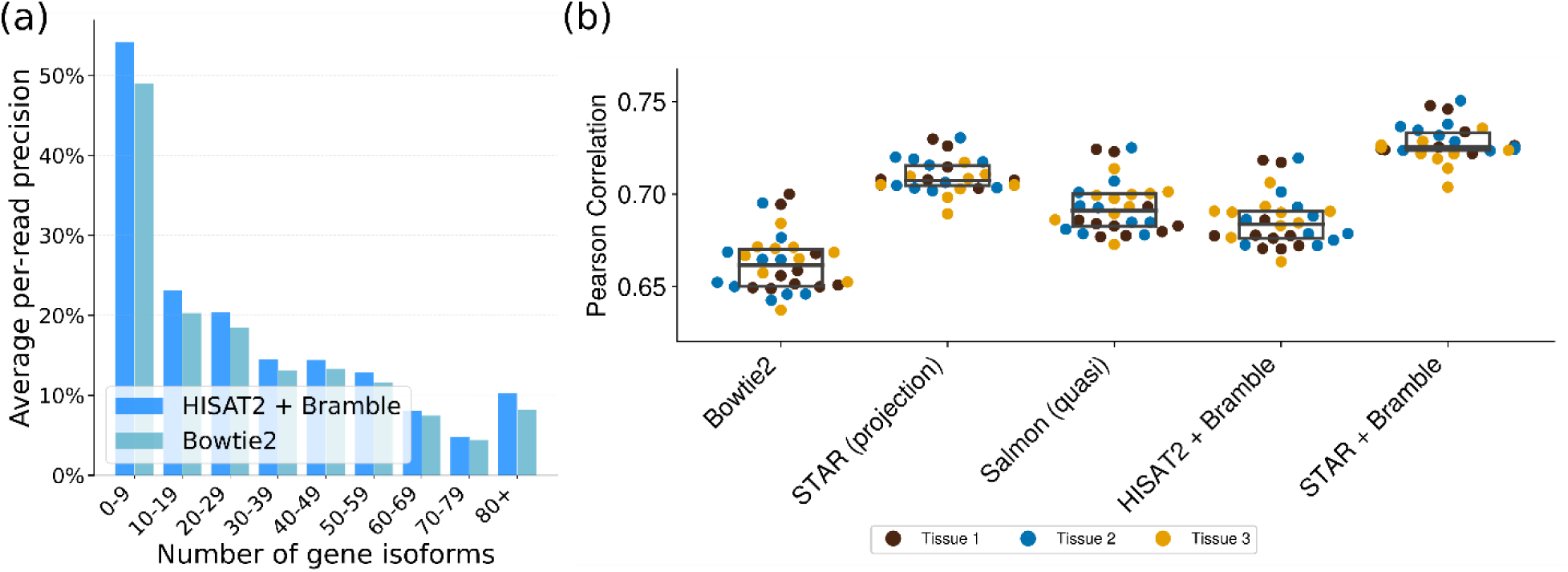
Performance of transcriptome alignment and transcript quantification pipelines on simulated short-read data. Results are averaged across three simulated tissue types with 10 samples each. a) Per-read precision for Bowtie2 run with -k 76 and HISAT + Bramble, averaged across reads and stratified by the number of annotated isoforms in the read’s gene of origin. For each read, precision was calculated as the number of correct alignments divided by the number of reported alignments. b) Swarm plot showing transcript quantification accuracy, measured by Pearson Correlation (PCC), for transcript abundances estimated from Bowtie2 run with -k 76, STAR transcriptome projection, HISAT2 + Bramble, STAR + Bramble, and Salmon quasi-mapping with decoys. For Bowtie2, STAR transcriptome projection, HISAT2 + Bramble, and STAR + Bramble, transcript abundances were estimated with Salmon in alignment-based mode using the corresponding transcriptomic alignments as input. Salmon quasi-mapping was run directly from reads. For boxes within swarms, center lines indicate medians, and lower and upper box edges indicate the first and third quartiles, respectively.

Next, we investigated whether the improved read-level precision affected downstream abundance estimates. We compared five short-read quantification pipelines (see Supplementary Note 1 for command line settings): Bowtie2 + Salmon, STAR (projection) + Salmon using STAR’s internal genome-to-transcriptome projection, Salmon quasi-mapping with decoys, HISAT2 + Bramble + Salmon, and STAR + Bramble + Salmon. We evaluated Pearson and Spearman correlation coefficients (PCC, SCC) and error metrics (MAE, RMSE, MAPE) between estimated and ground-truth counts, with metrics averaged across samples.

STAR + Bramble + Salmon performed best overall, achieving the highest PCC (0.728) and SCC (0.646) and the lowest MAE (110) and MAPE (64.2) (Fig. 1b; Supplementary Table 1; Supplementary Figs. 2-3). STAR + Bramble + Salmon was more similar in performance to STAR (projection) than to HISAT2 + Bramble + Salmon, suggesting that quantification accuracy depends strongly upon the underlying genomic alignments used for projection. HISAT2 + Bramble + Salmon performed better than Bowtie2 + Salmon in most metrics, with considerably higher PCC (0.686 vs. 0.663) and SCC (0.639 vs. 0.549). This suggests that improvements in transcriptomic alignment precision, as shown in Fig. 1a, may propagate to downstream transcript abundance estimates.

Transcripts with ground truth counts of zero but estimated abundances above zero were denoted false positive transcripts. Conversely, transcripts with ground truth counts above zero but estimated abundances of zero were denoted false negative transcripts. A substantial number of false positive transcripts were observed for every pipeline: HISAT2 + Bramble and Salmon (quasi) had the lowest numbers of false positives with low predicted abundance (<100 read counts) and STAR + Bramble + Salmon and STAR (projection) + Salmon had the lowest numbers of false positives with medium-to-high predicted abundance (≥100 read counts; Supplementary Fig. 4a). Fewer false negative transcripts were observed. Across all abundances, Salmon (quasi) had the lowest number of false negative transcripts (Supplementary Fig. 4b). Differences in the false positives and false negatives produced by each pipeline may contribute to the observed variation in correlation and error metrics between pipelines.

### Evaluation of performance on long-read data

We also benchmarked the performance of Bramble on long-read data using the simulated PacBio and ONT datasets and human reference annotation described in “Simulated data”. These datasets were simulated with tissue types, samples, and per-sample transcript abundances equivalent to those in the short-read dataset, and included the same added transcriptional noise. Long reads typically cover a substantial portion of their transcript of origin, allowing Bramble’s long-read mode to accurately project genomic alignments onto compatible annotated transcripts. We compared minimap2 genomic alignments projected to the transcriptome with Bramble against transcriptomic alignments produced by minimap2 (using parameters set up to report up to 116 distinct alignments; see “Secondary alignments in transcriptome alignment” for details). Supplementary Note 1 describes command line settings for each tool. We calculated precision, recall, and F1 score as described in the previous section and grouped reads by the number of isoforms belonging to their gene of origin.

For PacBio data, minimap2 + Bramble maintained relatively high precision (>50%) across bins, suggesting consistent performance regardless of gene locus complexity, while minimap2’s precision declined sharply as the complexity increased (Fig. 2a). Across bins, both methods had recall scores near 100% (Supplementary Fig. 5a); however, minimap2 + Bramble had higher F1 scores, with F1 scores approximately doubled compared to minimap2 for genes with more than 10 isoforms (Supplementary Fig. 5b). For ONT data, precision decreased with increasing locus complexity for both minimap2 + Bramble and minimap2 alone, but minimap2 + Bramble remained substantially more precise (Fig. 2b). Recall scores were lower than those observed for the PacBio data, potentially reflecting ONT-specific noise, and minimap2 alone had slightly higher recall (Supplementary Fig. 6a). However, minimap2 + Bramble yielded much higher F1 scores than minimap2 across every bin (Supplementary Fig. 6b).

**Figure 2.**
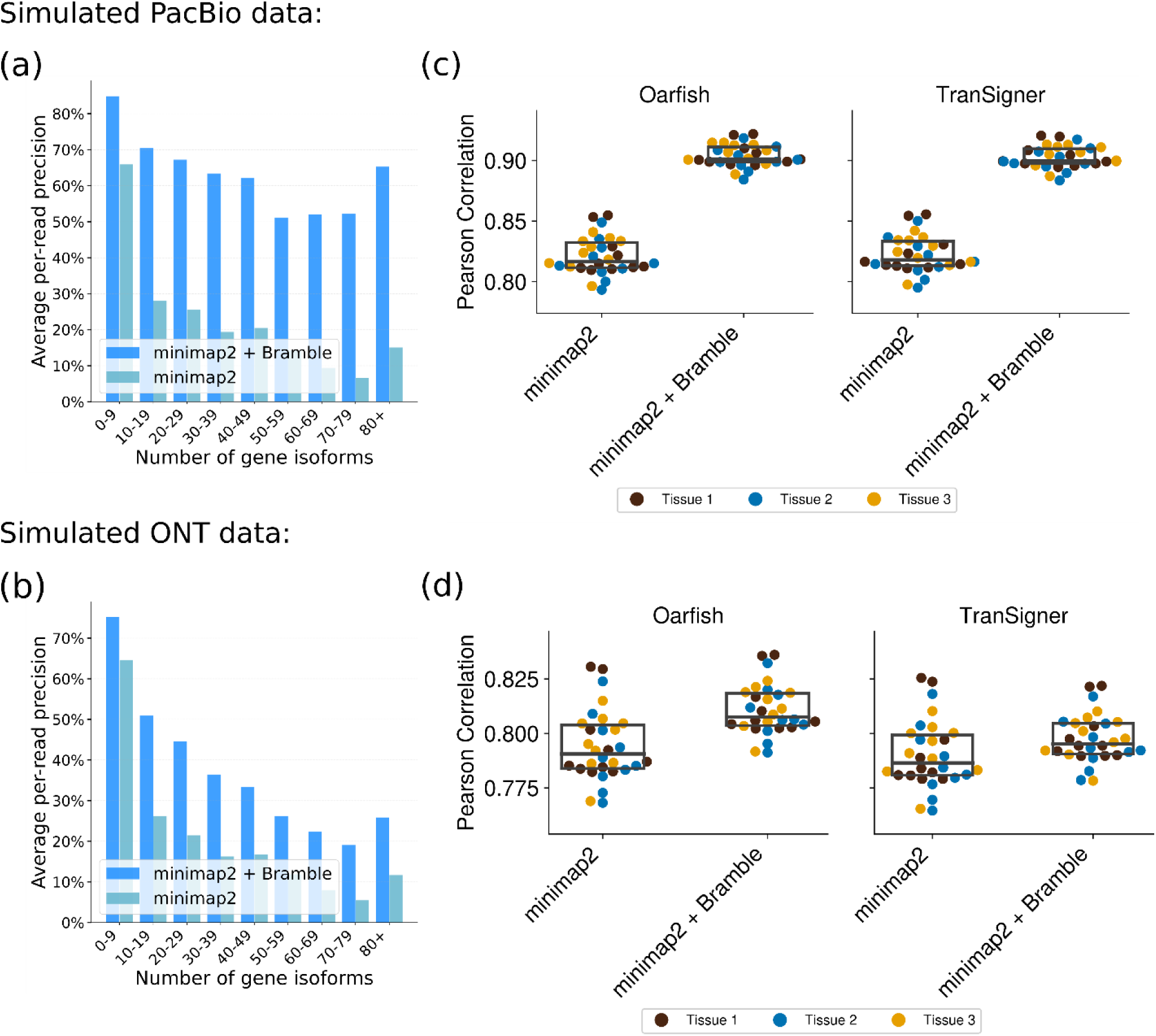
Performance of transcriptome alignment and transcript quantification pipelines on simulated long-read data. Results are averaged across three tissues and 10 samples per tissue. a) Per-read precision on simulated PacBio data, averaged across reads and stratified by the number of annotated isoforms in the read’s gene of origin. b) Transcript quantification accuracy on simulated PacBio data, measured by Pearson correlation coefficient (PCC). c) Per-read precision on simulated ONT data, averaged across reads and stratified by the number of annotated isoforms in the read’s gene of origin. d) Transcript quantification accuracy on simulated ONT data, measured by PCC. For a given read, precision is calculated as the number of correct alignments divided by the number of reported alignments. For boxes within swarms, center lines indicate medians and lower and upper box edges indicate the first and third quartiles, respectively.

After evaluating transcriptomic alignment accuracy across pipelines, we next evaluated downstream abundance estimates generated from the corresponding alignments. We used two long-read quantification methods, Oarfish and TranSigner, to test the consistency of results across approaches. We compared four quantification pipelines (see Supplementary Note 1 for command line settings): minimap2 + Oarfish, minimap2 + Bramble + Oarfish, minimap2 + TranSigner, and minimap2 + Bramble + TranSigner. For each pipeline, transcript abundances were estimated using the corresponding transcriptomic alignments as input, and performance was evaluated using Pearson and Spearman correlation coefficients and error metrics averaged across samples.

Bramble-based pipelines demonstrated substantial improvements in quantification accuracy, and these improvements were consistent across quantification methods. For both Oarfish and TranSigner on PacBio data, using Bramble considerably increased correlation coefficients (PCC: ∼0.82 to >0.90; SCC: 0.778 to 0.818; Fig. 2c; Supplementary Table 2; Supplementary Figs. 7-8), roughly halved MAE, RMSE, and MAPE (Supplementary Table 2; Supplementary Fig. 8), and reduced the number of false positive and false negative transcripts (Supplementary Fig. 9). Performance on ONT data showed a comparable pattern: for both quantification methods, using Bramble improved PCC, MAE, RMSE, and MAPE (Fig. 2d; Supplementary Table 3; Supplementary Figs. 10-11), and largely reduced the number of false positive and false negative transcripts (Supplementary Fig. 12). SCC decreased slightly, however, and improvements were more modest than for PacBio data, potentially due to ONT-specific noise.

### Evaluation of performance using incomplete reference annotations

The experiments in the previous sections used the human reference annotation, which is among the most complete and well-curated eukaryotic annotations available. However, many transcriptomic studies (e.g., studies on non-model organisms) use reference annotations that are far less complete. As annotation completeness decreases, annotation bias is expected to worsen: if a larger fraction of reads originates from transcripts absent from the reference, then there are more opportunities for read misassignment. We therefore benchmarked Bramble-based transcript quantification using incomplete reference annotations.

We subsampled the human reference annotation to retain 80%, 60%, 40%, and 20% of annotated transcripts, generating five replicate annotations at each retention level (see “Reduced annotations”). We evaluated performance on the same simulated RNA-seq data from the previous sections, using 10 samples from the first simulated tissue. We compared four quantification pipelines on short-read data: Bowtie2 + Salmon, Salmon quasi-mapping with decoys, HISAT2 + Bramble + Salmon, and STAR + Bramble + Salmon, and four quantification pipelines on long-read data: minimap2 + Oarfish, minimap2 + TranSigner, minimap2 + Bramble + Oarfish, and minimap2 + Bramble + TranSigner, using the command line settings described in Supplementary Note 1. Metrics were calculated using known simulated abundances as ground truth and averaged across samples and replicate annotations. Only transcripts retained in a given replicate annotation were included when comparing estimated and ground-truth abundances.

Using Bramble improved quantification accuracy across every read type and level of transcript retention, with lower transcript retention yielding larger improvements over baseline quantification pipelines. For short-read data, STAR + Bramble + Salmon consistently achieved the highest PCC values, followed by Salmon (quasi) at 80% transcript retention and HISAT2 + Bramble + Salmon at 20% to 60% transcript retention (Fig. 3a; Supplementary Tables 4-7; Supplementary Fig. 13). STAR + Bramble + Salmon also maintained the highest Spearman correlation and the lowest MAE, RMSE, and MAPE across retention levels (Supplementary Tables 4-7; Supplementary Fig. 14). For long-read data, Bramble-based pipelines generally outperformed their corresponding baseline pipelines, and the improvements were more pronounced than for short-read data. On PacBio data, minimap2 + Bramble + Oarfish and minimap2 + Bramble + TranSigner considerably outperformed minimap2 + Oarfish and minimap2 + TranSigner across all metrics (Fig. 3b; Supplementary Tables 8-11; Supplementary Figs. 15-16). Notably, for baseline pipelines, MAE, RMSE, and MAPE increased as transcript retention decreased, whereas for Bramble-based pipelines, these metrics remained nearly constant. Similar performance improvements were observed on ONT data, although SCC was lower at 80% transcript retention (Fig. 3c; Supplementary Tables 12-15; Supplementary Figs. 17-18). False positives and false negatives are reported for short-read, PacBio, and ONT data in Supplementary Figs. 19-24.

**Figure 3.**
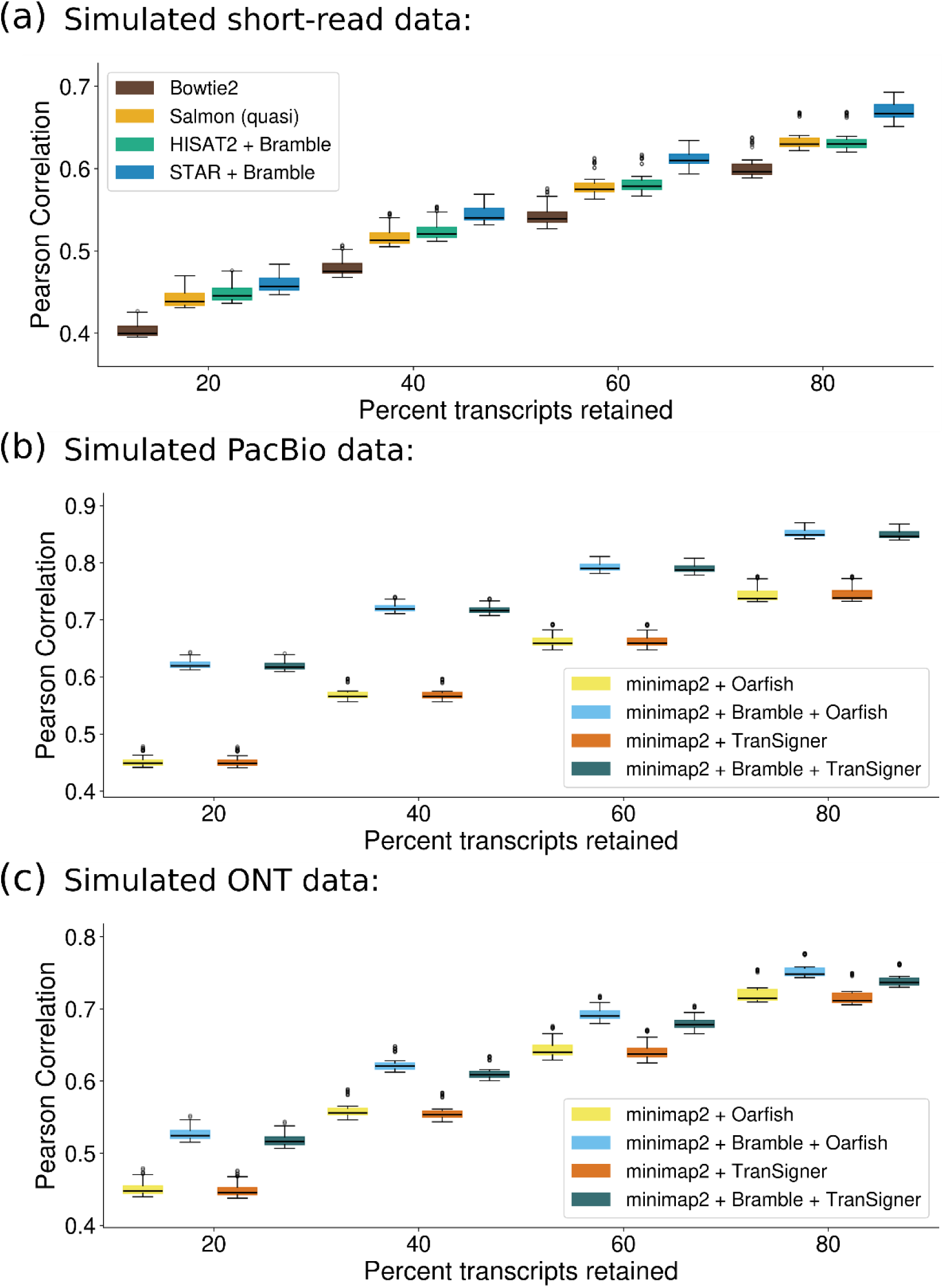
Performance of transcript quantification methods using subsampled human reference annotations with 20%, 40%, 60%, and 80% of transcripts retained. Results are averaged across 10 samples and five annotation replicates. Pearson correlation coefficient (PCC) on a) simulated short-read data, b) simulated PacBio data, and c) simulated ONT data. For box-and-whisker plots, center lines indicate medians and lower and upper box edges indicate the first and third quartiles, respectively.

## Discussion

Transcriptome-based quantification methods are limited by annotation bias, wherein reads originating from unannotated transcripts may be misassigned to annotated ones, skewing abundance estimates. We introduce Bramble, a software tool that improves transcript quantification by projecting spliced genomic alignments into transcriptomic space, reducing reliance on direct transcriptome mapping. Across short- and long-read data, Bramble-based quantification pipelines consistently produced more accurate transcript abundance estimates compared to conventional pipelines. Long-read data yielded particularly substantial improvements because long reads often span multiple splice junctions and cover a larger fraction of their transcript of origin, enabling Bramble to produce more accurate transcriptomic projections. Because annotation bias worsens when reference annotations are less complete, greater improvements were observed with simulated incomplete reference annotations, suggesting that Bramble is applicable across RNA-seq studies with varying levels of annotation completeness.

Bramble can be easily integrated into existing workflows; it is available as a standalone tool, as a direct integration within Salmon and Oarfish, and as a Rust library. Because it is compatible with any spliced genome aligner and any quantification method that accepts transcriptomic alignments, users can continue to work with their existing tools. Bramble can also simplify workflows: sequencing data are often stored in public repositories as genomic alignments, and Bramble can project these existing genomic alignments into transcriptomic coordinates for downstream quantification, eliminating the need to regenerate FASTQ files or realign reads with a transcriptome aligner.

While our benchmarks relied on a curated, fixed reference annotation, Bramble is also compatible with sample-specific transcript annotations generated by genome-guided assemblers such as StringTie^44^. Building a sample-specific annotation may better capture the cell types represented in the sample and improve quantification accuracy. Because genome-guided transcript assembly already requires genomic alignments, applying Bramble to the same alignments is straightforward. In such cases, Bramble allows the same alignments to serve for quantification as well, reducing the computation and I/O burden that results from having to perform redundant transcriptome alignment. The same approach can be used for non-model or newly sequenced organisms with limited existing annotation.

## Methods

### Algorithm

### Building the genome-to-transcriptome tree

Bramble uses the reference annotation to build an index over the genomic intervals spanned by annotated transcript exons. Let the reference annotation define a set of transcripts *T* = {*t*_1_,…, *t_n_*}, where each transcript *t_i_* is an ordered sequence of exons on a fixed chromosome and strand *t_i_* = (*e_i_*_,1_, *e_i_*_,2_,…, *e_i_*_,*n*_*_i_*), and each exon is a genomic interval *e_i_*_,*l*_ = [*u_i_*_,*l*_, *v_i_*_,*l*_].

For every chromosome-strand pair (*c*, *σ*), Bramble builds an implicit interval tree^45^ *I_c_*_,*σ*_ over the exon intervals associated with that pair, *E_c_*_,*σ*_ = {*e_i_*_,*l*_: *t_i_* is on (*c*, *σ*), 1 ≤ *l* ≤ *n_i_*}. We denote *G*(*c*, *σ*) = *I_c_*,*_σ_* the tree associated with a chromosome *c* and strand *σ*, and refer to the full collection *G* = {*I_c_*,*_σ_*}*_c_*,*_σ_* as the genome-to-transcriptome tree, or g2t tree (Fig. 4a). Together, these trees index the full reference exon set *E* = ⋃*_c_*_,*σ*_ *E_c_*_,*σ*_. The trees are built once at program start in *O*(*N* log*N*) time, where *N* = |*E*| is the total number of reference exons.

**Figure 4.**
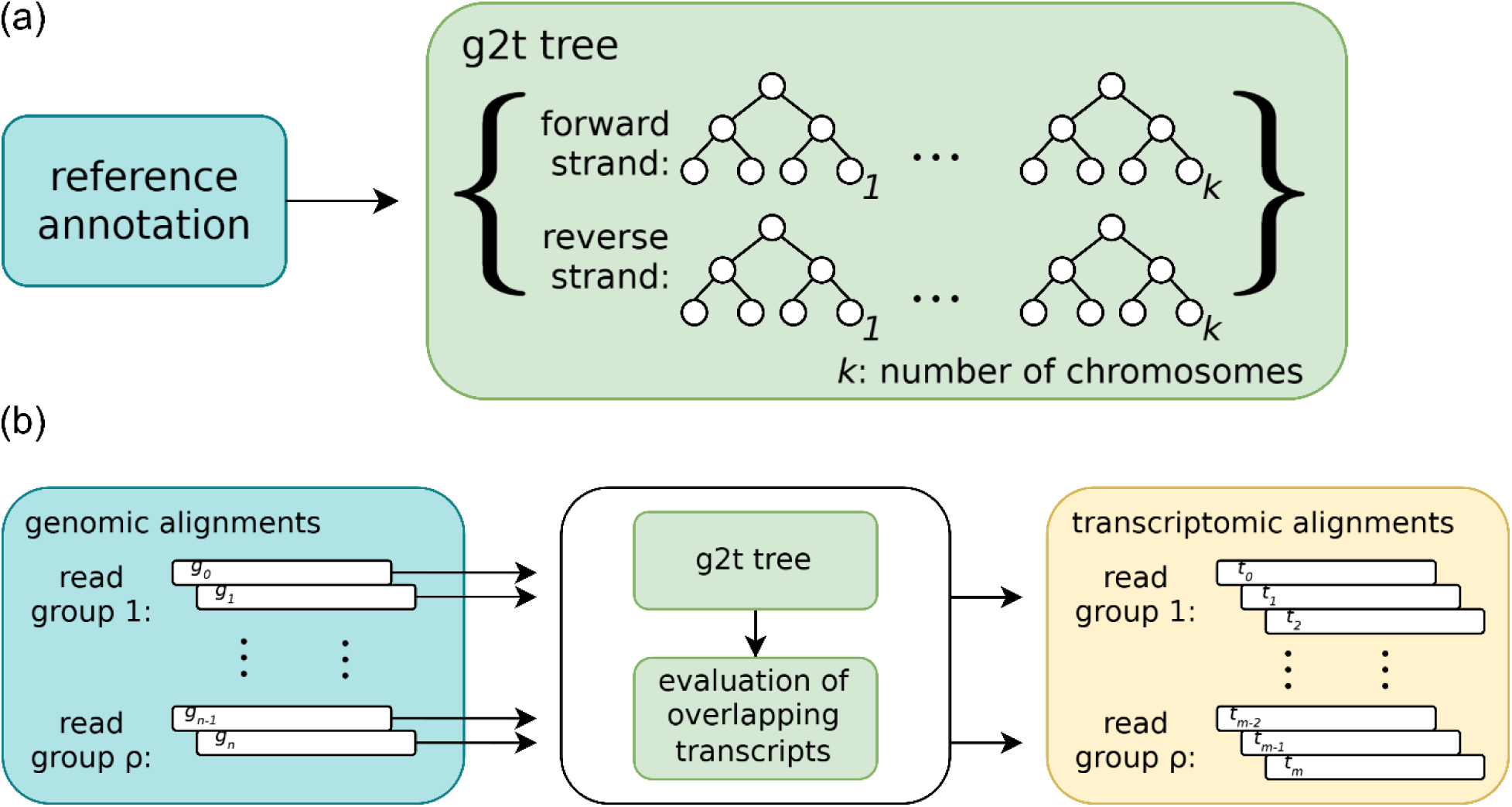
Construction and use of Bramble’s genome-to-transcriptome (g2t) tree. a) The reference annotation is used to build the g2t tree, an index of genomic intervals corresponding to annotated transcript exons. The g2t tree is implemented as a collection of implicit interval trees^45^, with one tree per chromosome and strand. b) Bramble uses the g2t tree to project genomic alignments into transcriptomic coordinates. Genomic alignments are processed by read, with each read group representing all genomic alignments reported for a single sequencing read. For each genomic alignment, the g2t tree identifies overlapping annotated transcripts, and an evaluation step determines which candidate transcripts are compatible with the alignment. A read group may contain one or more genomic alignments and may produce zero, one, or multiple transcriptomic alignments.

In addition to interval coordinates, each tree node stores metadata identifying the reference transcript to which the exon belongs. Because the same genomic interval can be shared by multiple transcripts, intervals with identical genomic coordinates are stored independently in the tree rather than collapsed into a single node.

### Querying the genome-to-transcriptome tree

Bramble uses the genome-to-transcriptome tree to identify transcriptomic alignments compatible with a given genomic alignment (Fig. 4b). We represent each genomic alignment by its exon chain, ordered by genomic coordinate. Thus, a genomic alignment *A* is written as *A* = (*a*_1_,…, *a_p_*), where each aligned exon is a genomic interval *a_j_* = [*α_j_*, *β_j_*).

Each aligned exon *a_j_* is queried against *G* to find the transcripts it could have plausibly originated from. Given the chromosome and strand (*c*, *σ*) of *a_j_*, Bramble retrieves the corresponding tree *G*(*c*, *σ*) = *I_c_*,*_σ_* and queries it for reference exons overlapping *a_j_*, together with the transcript to which each exon belongs:

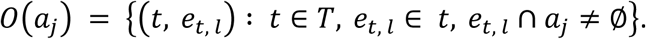

This query runs in *O*(log*N_c_*,*_σ_* + *m*) time, where *N_c_*,*_σ_* = |*E_c_*,*_σ_*| is the number of reference exon intervals in the queried chromosome-strand tree and *m* = |*O*(*a_j_*)| is the number of overlapping exons returned.

Each candidate pair in *O*(*a_j_*) is then filtered using positional offset thresholds at the exon boundaries to rule out transcripts whose exon structure is inconsistent with the alignment. For a candidate transcript-exon pair (*t*, *e_t_*_,*l*_) ∈ *O*(*a_j_*), with *a_j_* = [*α_j_*, *β_j_*) and *e_t_*_,*l*_ = [*u_t_*_,*l*_, *v_t_*_,*l*_), define the left and right boundary offsets as

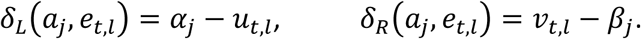

Each side *x* ∈ {*L*, *R*} is classified along two independent axes. First, by the sign of *δ_x_*: *δ_x_* < 0 indicates that the alignment extends past the reference exon boundary on that side, which is treated as an insertion of size |*δ_x_*| relative to the transcript; *δ_x_* > 0 indicates that the reference exon extends past the alignment boundary, which is treated as a deletion of size *δ_x_* relative to the alignment; and *δ_x_* = 0 indicates an exact boundary match. Second, each side is classified by its position in the exon chain: the left side of *a*_1_ and the right side of *a_p_* are *terminal*, marking the ends of the aligned region, whereas all other sides are *internal*.

Whether a candidate pair (*t*, *e_t_*_,*l*_) passes filtering at side *x* depends upon both classifications. At a terminal side, a deletion (*δ_x_* > 0) passes unconditionally, because a read is not expected to cover the full annotated exon or transcript boundary. For instance, a short read may cover only one or two exons of a much longer transcript. An insertion (*δ_x_* < 0) indicates that the alignment extends past the reference exon into adjacent intronic sequence; this can be reconciled with the transcript model only if the overhang is small enough to be soft clipped, so it passes only if |*δ_x_*| ≤ *τ_clip_*. At an internal side, a deletion relative to the transcript model passes if *δ_x_* ≤ *τ_del_*, and an insertion relative to the transcript model passes if |*δ_x_*| ≤ *τ_ins_*. Exact matches (*δ_x_* = 0) pass trivially.

A candidate pair (*t*, *e_t_*_,*l*_) is retained only if it passes filtering at both the left (*L*) and right (*R*) boundaries of *a_j_*. After filtering, the result of querying *G* with *a_j_* is

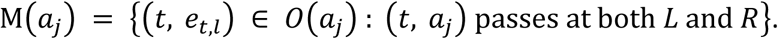

### Resolving alignment-level matches

A transcript *t* is a valid projection target only if the alignment exons correspond one-to-one with the exons of *t* lying within the aligned region: every alignment exon maps to exactly one transcript exon, and every transcript exon between the first and last matched positions maps to exactly one alignment exon. Since an alignment is not expected to reach either end of the transcript it originated from, exons of *t* outside this region are unconstrained.

Because the exons of *t* are non-overlapping and *a_j_* is a single genomic interval, *a_j_* can match at most one exon of *t*. For *t* ∈ *T*, let *l_j_*(*t*) denote the index of the exon matched to *a_j_* in *M*(*a_j_*), when one such match exists. Define the transcript-level candidate set for *a_j_* as

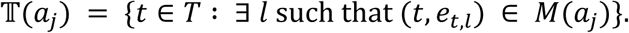

The alignment-level set of matched transcripts is then

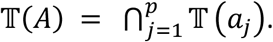

If some *a_j_* has no matching transcript exon, T(*a_j_*) = ∅, and therefore T(*A*) = ∅.

Intersection alone does not guarantee that matched exons are consecutive in *t*: a transcript could contain an extra exon between the positions matched to *a_j_* and *a_j_*_+1_ and still appear in both T(*a_j_*) and T(*a_j_* _+ 1_). We therefore additionally require

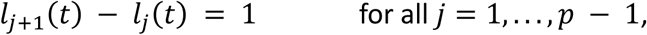

and remove from T(*A*) any transcript that fails this condition for any *j*.

### Reporting transcriptomic alignments

For every transcript in T(*A*), alignment information is compiled, including the alignment position, CIGAR string, and alignment score. Supplementary Note 2 describes Bramble’s implementation of clip-rescue, which may be used with long-read data to improve the discriminatory power of alignment scores; Supplementary Note 3 describes Bramble’s formula for calculating alignment score.

If the genomic alignment *A* comes from an unpaired read, then corresponding transcriptomic alignments are reported for every transcript in T(*A*). If there are multiple genomic alignments for the same read, then the transcriptomic alignments generated using all genomic alignments are compiled together before they are reported, so that metrics including mapping quality (see Supplementary Note 4) can be computed. For paired-end data, Bramble applies filtering rules to ensure consistency between the transcriptomic alignments reported by the mates in each mate pair. Supplementary Note 5 describes these rules.

### Simulated data

To evaluate the performance of Bramble-based quantification pipelines, we used short- and long-read simulated data. For short-read data, we used the dataset from Varabyou et al. (2021), which includes realistic transcriptional noise. In that simulation, Polyester^46^ was used to generate forward-stranded reads from the GRCh38 reference genome sequence and an expanded version of the CHESS 2.2 reference annotation^34, 22^. The dataset contains three simulated tissue types, each with ten samples. To randomize read order, we shuffled reads using seqkit shuffle^47^.

For long-read data, we used SQANTI-SIM^48^ to simulate ONT and PacBio reads with added transcriptional noise. The ONT and PacBio datasets each contain three simulated tissue types with ten samples per tissue, modeled after the short-read dataset: we simulated known and novel transcripts for each sample using the TPM values of the transcript in the corresponding short-read sample.

### Secondary alignments in transcriptome alignment

Accurate quantification requires reporting complete sets of secondary alignments for multi-mapped reads, because the expectation-maximization (EM) algorithms used by quantification tools estimate isoform abundances by weighting each read across every compatible transcript, so incomplete sets of alignments bias those estimates towards the isoforms that happen to be reported^5, 9^. Computing and reporting every possible alignment, however, carries substantial computational cost, particularly for short reads, which multi-map more often, frequently matching many isoforms that share exons.

For our benchmarks, we configured Bowtie2 and minimap2 to report a near-complete set of secondary alignments. To estimate an upper bound on the number of possible secondary alignments at any locus, we counted overlapping transcripts per gene locus and found a maximum of 116 in the expanded CHESS 2.2 annotation: at most one primary alignment plus 115 secondary alignments for a read compatible with all transcripts at that locus (Supplementary Fig. 25 shows linear- and log-scale histograms of overlapping transcripts per gene locus). Therefore, for transcriptome alignment, we used -N 115 for minimap2 and-k 76 for Bowtie2, which was the highest value at which Salmon’s alignment-based mode ran reliably; higher values made Bowtie2’s output crash Salmon.

### Reduced annotations

We created reduced annotations by randomly subsampling the expanded CHESS 2.2 annotation at the transcript-level, retaining 80%, 60%, 40%, and 20% of annotated transcripts. We selected entries for retention by uniform random sampling without replacement. Because genes remained represented in an annotation as long as they retained at least one transcript, the proportion of retained genes exceeded the proportion of retained transcripts at every retention level. To reduce sensitivity to the specific genes or transcripts sampled, we generated five random subsampling replicates of each reduced annotation at every retention level, each initialized with a pre-determined seed.

### Versions

The following software versions were used: HISAT2 version 2.2.1, STAR version 2.7.11b, Bowtie2 version 2.5.4, Salmon version 1.10.3, minimap2 version 2.30-r1287, Oarfish version 0.9.0, TranSigner version 1.2.0, and Samtools version 1.22.1.

## Data availability

The simulated short- and long-read datasets are available at ftp://ftp.ccb.jhu.edu/pub/data/bramble. The reference genome, annotation, and transcriptome used in this study are available at the same location. Scripts for benchmarking and data analysis, as well as comprehensive documentation describing the datasets and scripts, are provided at https://github.com/zrudnick/bramble-scripts.

## Code availability

Bramble is an open-source software provided on GitHub (https://github.com/zrudnick/bramble), which is the recommended avenue for submission of issues or suggestions. C++ and Rust implementations are provided. Bramble is also available on Bioconda.

Bramble is provided under an MIT license.

## Acknowledgments

This work was supported in part by NSF grant DBI-2412449 (M.P.), NIH grant R35-GM156470 (M.P.), and NIH grant R01HG009937 (R.P.).

## Author contributions

Z.R., A.V., M.P. conceived the study. Z.R., A.V., and R.P. developed Bramble, and Z.R. and A.V. performed the benchmarking analyses. Z.R. wrote the manuscript with input from all authors. R.P. and M.P. supervised the project.

## References

1. Langmead, B., Trapnell, C., Pop, M. & Salzberg, S. L. Ultrafast and memory-efficient alignment of short DNA sequences to the human genome. Genome Biol 10, R25 (2009).

2. Li, H. & Durbin, R. Fast and accurate short read alignment with Burrows–Wheeler transform. Bioinformatics 25, 1754–1760 (2009).

3. Langmead, B. & Salzberg, S. L. Fast gapped-read alignment with Bowtie 2. Nat Methods 9, 357–359 (2012).

4. Li, H. Aligning sequence reads, clone sequences and assembly contigs with BWA-MEM. (2013).

5. Li, B. & Dewey, C. N. RSEM: accurate transcript quantification from RNA-Seq data with or without a reference genome. BMC Bioinformatics 12, 323 (2011).

6. Patro, R., Mount, S. M. & Kingsford, C. Sailfish enables alignment-free isoform quantification from RNA-seq reads using lightweight algorithms. Nat. Biotechnol. 32, 462–464 (2014).

7. Zhang, Z. & Wang, W. RNA-Skim: a rapid method for RNA-Seq quantification at transcript level. Bioinformatics 30, i283–i292 (2014).

8. Bray, N. L., Pimentel, H., Melsted, P. & Pachter, L. Near-optimal probabilistic RNA-seq quantification. Nat Biotechnol 34, 525–527 (2016).

9. Patro, R., Duggal, G., Love, M. I., Irizarry, R. A. & Kingsford, C. Salmon provides fast and bias-aware quantification of transcript expression. Nat Methods 14, 417–419 (2017).

10. Sarkar, H., Zakeri, M., Malik, L. & Patro, R. Towards Selective-Alignment: Bridging the Accuracy Gap between Alignment-Based and Alignment-Free Transcript Quantification (Proceedings of the 2018 ACM International Conference on Bioinformatics, Computational Biology, and Health Informatics, Association for Computing Machinery, New York, NY, USA, 2018).

11. Srivastava, A., et al. Alignment and mapping methodology influence transcript abundance estimation. Genome Biol. 21, 239–8 (2020).

12. Wang, Y., Zhao, Y., Bollas, A., Wang, Y. & Au, K. F. Nanopore sequencing technology, bioinformatics and applications. Nat. Biotechnol. 39, 1348–1365 (2021).

13. Wongsurawat, T., Jenjaroenpun, P. & Nookaew, I. Direct Sequencing of RNA and RNA Modification Identification Using Nanopore. Methods Mol. Biol. 2477, 71–77 (2022).

14. Rhoads, A. & Au, K. F. PacBio Sequencing and Its Applications. Genomics Proteomics Bioinformatics 13, 278–289 (2015).

15. Wenger, A. M., et al. Accurate circular consensus long-read sequencing improves variant detection and assembly of a human genome. Nat. Biotechnol. 37, 1155–1162 (2019).

16. Li, H. Minimap2: pairwise alignment for nucleotide sequences. Bioinformatics 34, 3094–3100 (2018).

17. Zare Jousheghani, Z., Singh, N. P. & Patro, R. Oarfish: enhanced probabilistic modeling leads to improved accuracy in long read transcriptome quantification. Bioinformatics 41(Supplement_1), i304–i313 (2025).

18. Ji, H. J. & Pertea, M. Enhancing transcriptome expression quantification through accurate assignment of long RNA sequencing reads with TranSigner. Genome Biol 26, 257 (2025).

19. Loving, R. K., et al. Long-read sequencing transcriptome quantification with lr-kallisto. PLoS Comput Biol 21, e1013692 (2025).

20. Ma, C. & Kingsford, C. Detecting, Categorizing, and Correcting Coverage Anomalies of RNA-Seq Quantification. Cell. Syst. 9, 589–599.e7 (2019).

21. Chisanga, D., Liao, Y. & Shi, W. Impact of gene annotation choice on the quantification of RNA-seq data. BMC Bioinformatics 23, 107 (2022).

22. Varabyou, A., et al. CHESS 3: an improved, comprehensive catalog of human genes and transcripts based on large-scale expression data, phylogenetic analysis, and protein structure. Genome Biol 24, 249 (2023).

23. Amaral, P., et al. The status of the human gene catalogue. Nature 622, 41–47 (2023).

24. Zhang, Q. & Shao, M. Transcript assembly and annotations: Bias and adjustment. PLoS Comput Biol 19, e1011734 (2023).

25. Ji, H. J., Pertea, M. & Salzberg, S. L. Annotating genomes at increased scale and resolution. Nature Reviews Genetics (2026).

26. Alvarez, M., Schrey, A. W. & Richards, C. L. Ten years of transcriptomics in wild populations: what have we learned about their ecology and evolution? Mol. Ecol. 24, 710–725 (2015).

27. Vuruputoor, V. S., et al. Welcome to the big leaves: Best practices for improving genome annotation in non-model plant genomes. Appl. Plant Sci 11, e11533 (2023).

28. Struhl, K. Transcriptional noise and the fidelity of initiation by RNA polymerase II. Nat Struct Mol Biol 14, 103–105 (2007).

29. Clark, M. B., et al. The Reality of Pervasive Transcription. PLOS Biology 9, e1000625 (2011).

30. Djebali, S., et al. Landscape of transcription in human cells. Nature 489, 101–108 (2012).

31. Palazzo, A. F. & Lee, E. S. Non-coding RNA: what is functional and what is junk? Front. Genet. 6, 2 (2015).

32. Cavallaro, M., et al. 3 ′-5 ′ crosstalk contributes to transcriptional bursting. Genome Biol 22, 56 (2021).

33. Wan, Y., et al. Dynamic imaging of nascent RNA reveals general principles of transcription dynamics and stochastic splice site selection. Cell 184, 2878–2895.e20 (2021).

34. Pertea, M., et al. CHESS: a new human gene catalog curated from thousands of large-scale RNA sequencing experiments reveals extensive transcriptional noise. Genome Biol 19 (2018).

35. Varabyou, A., Salzberg, S. L. & Pertea, M. Effects of transcriptional noise on estimates of gene and transcript expression in RNA sequencing experiments. Genome Res. 31, 301–308 (2021).

36. Pavey, S. A., Bernatchez, L., Aubin-Horth, N. & Landry, C. R. What is needed for next-generation ecological and evolutionary genomics? Trends in Ecology & Evolution 27, 673–678 (2012).

37. Sundaram, A., Tengs, T. & Grimholt, U. Issues with RNA-seq analysis in non-model organisms: A salmonid example. Developmental & Comparative Immunology 75, 38–47 (2017).

38. Varabyou, A., et al. Comprehensive Transcriptome Annotation of Thousands of HIV-1 Genomes. bioRxiv (2025).

39. Roberts, S. A. & Gordenin, D. A. Hypermutation in human cancer genomes: footprints and mechanisms. Nature Reviews Cancer 14, 786–800 (2014).

40. Watson, C. T., et al. Complete Haplotype Sequence of the Human Immunoglobulin Heavy-Chain Variable, Diversity, and Joining Genes and Characterization of Allelic and Copy-Number Variation. The American Journal of Human Genetics 92, 530–546 (2013).

41. Deschamps-Francoeur, G., Simoneau, J. & Scott, M. S. Handling multi-mapped reads in RNA-seq. Computational and Structural Biotechnology Journal 18, 1569–1576 (2020).

42. Dobin, A., et al. STAR: ultrafast universal RNA-seq aligner. Bioinformatics 29, 15–21 (2012).

43. Kim, D., Paggi, J. M., Park, C., Bennett, C. & Salzberg, S. L. Graph-based genome alignment and genotyping with HISAT2 and HISAT-genotype. Nat. Biotechnol. 37, 907–915 (2019).

44. Pertea, M., et al. StringTie enables improved reconstruction of a transcriptome from RNA-seq reads. Nat Biotechnol 33, 290–295 (2015).

45. Li, H. & Rong, J. Bedtk: finding interval overlap with implicit interval tree. Bioinformatics 37, 1315–1316 (2021).

46. Frazee, A. C., Jaffe, A. E., Langmead, B. & Leek, J. T. Polyester: simulating RNA-seq datasets with differential transcript expression. Bioinformatics 31, 2778–2784 (2015).

47. Shen, W., Sipos, B. & Zhao, L. SeqKit2: A Swiss army knife for sequence and alignment processing. iMeta 3 (2024).

48. Mestre-Tomás, J., Liu, T., Pardo-Palacios, F. & Conesa, A. SQANTI-SIM: a simulator of controlled transcript novelty for lrRNA-seq benchmark. Genome Biol 24, 286 (2023).

49. Pertea, G. & Pertea, M. GFF Utilities: GffRead and GffCompare. F1000Res 9, 10.12688/f1000research.23297.2. eCollection 2020 (2020).

50. Suzuki, H. & Kasahara, M. Introducing difference recurrence relations for faster semi-global alignment of long sequences. BMC Bioinformatics 19, 45 (2018).

51. Kim, D., et al. TopHat2: accurate alignment of transcriptomes in the presence of insertions, deletions and gene fusions. Genome Biol 14, R36 (2013).

