## Supplementary Material for "Bramble: projection of spliced genomic alignments into transcriptomic space for improved transcript quantification"

| Method | PCC | SCC | MAE | RMSE | MAPE |
|---|---|---|---|---|---|
| HISAT2 + Bramble + Salmon | 0.686 | 0.639 | 129 | 3100 | 76.5 |
| STAR + Bramble + Salmon | **0.728** | **0.646** | **110** | 3270 | **64.2** |
| STAR (projection) + Salmon | 0.710 | 0.610 | 111 | **2990** | 65.1 |
| Bowtie + Salmon | 0.663 | 0.549 | 130 | 3290 | 76.2 |
| Salmon (quasi) | 0.693 | 0.645 | 126 | 3430 | 71.9 |

**Supplementary Table 1.** Correlation and error metrics for simulated short-read data.

| Method | PCC | SCC | MAE | RMSE | MAPE |
|---|---|---|---|---|---|
| minimap2 + Bramble + Oarfish | **0.904** | **0.818** | **3.76** | **237** | **18.8** |
| minimap2 + Oarfish | 0.821 | 0.778 | 8.79 | 522 | 48.5 |
| minimap2 + Bramble + TranSigner | 0.903 | **0.818** | 3.77 | 238 | 22.4 |
| minimap2 + TranSigner | 0.823 | 0.778 | 8.67 | 523 | 50.6 |

**Supplementary Table 2.** Correlation and error metrics for simulated PacBio data.

| Method | PCC | SCC | MAE | RMSE | MAPE |
|---|---|---|---|---|---|
| minimap2 + Bramble + Oarfish | **0.811** | 0.680 | **7.11** | **255** | **50.9** |
| minimap2 + Oarfish | 0.794 | **0.721** | 9.47 | 493 | 63.1 |
| minimap2 + Bramble + TranSigner | 0.798 | 0.676 | 7.53 | 256 | 51.5 |
| minimap2 + TranSigner | 0.790 | 0.719 | 9.59 | 498 | 64.5 |

**Supplementary Table 3.** Correlation and error metrics for simulated ONT data.

| Method | PCC | SCC | MAE | RMSE | MAPE |
|---|---|---|---|---|---|
| HISAT2 + Bramble + Salmon | 0.450 | 0.429 | 796 | 7660 | 439 |
| STAR + Bramble + Salmon | **0.461** | **0.434** | **668** | **7220** | **357** |
| Bowtie + Salmon | 0.404 | 0.367 | 850 | 8020 | 499 |
| Salmon (quasi) | 0.443 | 0.420 | 815 | 7910 | 455 |

**Supplementary Table 4.** Correlation and error metrics for simulated short-read data with reduced annotation comprising 20% transcripts from reference genome annotation.

| Method | PCC | SCC | MAE | RMSE | MAPE |
|---|---|---|---|---|---|
| HISAT2 + Bramble + Salmon | 0.526 | 0.498 | 424 | 6590 | 213 |
| STAR + Bramble + Salmon | **0.545** | **0.503** | **347** | **6230** | **163** |
| Bowtie + Salmon | 0.481 | 0.420 | 445 | 6700 | 227 |
| Salmon (quasi) | 0.519 | 0.487 | 431 | 6580 | 213 |

**Supplementary Table 5.** Correlation and error metrics for simulated short-read data with reduced annotation comprising 40% transcripts from reference genome annotation.

| Method | PCC | SCC | MAE | RMSE | MAPE |
|---|---|---|---|---|---|
| HISAT2 + Bramble + Salmon | 0.584 | 0.550 | 259 | 3900 | 147 |
| STAR + Bramble + Salmon | **0.612** | **0.557** | **210** | **3710** | **113** |
| Bowtie + Salmon | 0.544 | 0.463 | 269 | 4090 | 152 |
| Salmon (quasi) | 0.581 | 0.543 | 260 | 3950 | 148 |

**Supplementary Table 6.** Correlation and error metrics for simulated short-read data with reduced annotation comprising 60% transcripts from reference genome annotation.

| Method | PCC | SCC | MAE | RMSE | MAPE |
|---|---|---|---|---|---|
| HISAT2 + Bramble + Salmon | 0.636 | 0.596 | 183 | 3730 | 108 |
| STAR + Bramble + Salmon | **0.670** | **0.602** | **150** | **3560** | **84.5** |
| Bowtie + Salmon | 0.603 | 0.505 | 187 | 3890 | 108 |
| Salmon (quasi) | 0.637 | 0.594 | 181 | 3780 | 103 |

**Supplementary Table 7.** Correlation and error metrics for simulated short-read data with reduced annotation comprising 80% transcripts from reference genome annotation.

| Method | PCC | SCC | MAE | RMSE | MAPE |
|---|---|---|---|---|---|
| minimap2 + Bramble + Oarfish | **0.622** | 0.541 | **23.7** | **404** | **120** |
| minimap2 + Oarfish | 0.453 | 0.432 | 117 | 2940 | 988 |
| minimap2 + Bramble + TranSigner | 0.620 | **0.543** | **23.7** | 408 | **120** |
| minimap2 + TranSigner | 0.453 | 0.433 | 117 | 2890 | 992 |

**Supplementary Table 8.** Correlation and error metrics for simulated PacBio data with reduced annotation comprising 20% transcripts from reference genome annotation.

| Method | PCC | SCC | MAE | RMSE | MAPE |
|---|---|---|---|---|---|
| minimap2 + Bramble + Oarfish | **0.722** | 0.626 | **15.4** | **694** | **74.1** |
| minimap2 + Oarfish | 0.571 | 0.538 | 51.4 | 2060 | 259 |
| minimap2 + Bramble + TranSigner | 0.719 | **0.628** | **15.4** | 701 | 76.5 |
| minimap2 + TranSigner | 0.571 | 0.539 | 51.3 | 2050 | 259 |

**Supplementary Table 9.** Correlation and error metrics for simulated PacBio data with reduced annotation comprising 40% transcripts from reference genome annotation.

| Method | PCC | SCC | MAE | RMSE | MAPE |
|---|---|---|---|---|---|
| minimap2 + Bramble + Oarfish | **0.793** | 0.696 | 9.02 | **275** | **48.8** |
| minimap2 + Oarfish | 0.664 | 0.624 | 26.1 | 920 | 142 |
| minimap2 + Bramble + TranSigner | 0.790 | **0.697** | **9.01** | 278 | 54.6 |
| minimap2 + TranSigner | 0.664 | 0.624 | 26.0 | 918 | 150 |

**Supplementary Table 10.** Correlation and error metrics for simulated PacBio data with reduced annotation comprising 60% transcripts from reference genome annotation.

| Method | PCC | SCC | MAE | RMSE | MAPE |
|---|---|---|---|---|---|
| minimap2 + Bramble + Oarfish | **0.852** | **0.759** | **5.75** | **216** | **37.9** |
| minimap2 + Oarfish | 0.746 | 0.703 | 15.5 | 688 | 94.0 |
| minimap2 + Bramble + TranSigner | 0.850 | **0.759** | **5.75** | 221 | 43.8 |
| minimap2 + TranSigner | 0.746 | 0.703 | 15.3 | 737 | 99.7 |

**Supplementary Table 11.** Correlation and error metrics for simulated PacBio data with reduced annotation comprising 80% transcripts from reference genome annotation.

| Method | PCC | SCC | MAE | RMSE | MAPE |
|---|---|---|---|---|---|
| minimap2 + Bramble + Oarfish | **0.528** | **0.469** | **44.8** | 699 | **251** |
| minimap2 + Oarfish | 0.452 | 0.419 | 103 | 2020 | 682 |
| minimap2 + Bramble + TranSigner | 0.519 | 0.466 | 44.9 | **684** | 254 |
| minimap2 + TranSigner | 0.450 | 0.418 | 103 | 2010 | 679 |

**Supplementary Table 12.** Correlation and error metrics for simulated ONT data with reduced annotation comprising 20% transcripts from reference genome annotation.

| Method | PCC | SCC | MAE | RMSE | MAPE |
|---|---|---|---|---|---|
| minimap2 + Bramble + Oarfish | **0.624** | **0.539** | **26.7** | **1160** | **146** |
| minimap2 + Oarfish | 0.561 | 0.510 | 48.6 | 1730 | 255 |
| minimap2 + Bramble + TranSigner | 0.613 | 0.536 | 26.9 | 1200 | 151 |
| minimap2 + TranSigner | 0.558 | 0.510 | 48.7 | 1790 | 257 |

**Supplementary Table 13.** Correlation and error metrics for simulated ONT data with reduced annotation comprising 40% transcripts from reference genome annotation.

| Method | PCC | SCC | MAE | RMSE | MAPE |
|---|---|---|---|---|---|
| minimap2 + Bramble + Oarfish | **0.694** | **0.592** | **15.1** | 306 | **100** |
| minimap2 + Oarfish | 0.646 | 0.584 | 25.6 | 716 | 165 |
| minimap2 + Bramble + TranSigner | 0.681 | 0.587 | 15.4 | **289** | 105 |
| minimap2 + TranSigner | 0.642 | 0.583 | 25.7 | 720 | 172 |

**Supplementary Table 14.** Correlation and error metrics for simulated ONT data with reduced annotation comprising 60% transcripts from reference genome annotation.

| Method | PCC | SCC | MAE | RMSE | MAPE |
|---|---|---|---|---|---|
| minimap2 + Bramble + Oarfish | **0.754** | **0.637** | **10.1** | 249 | **73.8** |
| minimap2 + Oarfish | 0.723 | 0.654 | 15.1 | 546 | 103 |
| minimap2 + Bramble + TranSigner | 0.741 | 0.632 | 10.4 | **233** | 77.8 |
| minimap2 + TranSigner | 0.719 | 0.652 | 15.2 | 544 | 107 |

**Supplementary Table 15.** Correlation and error metrics for simulated ONT data with reduced annotation comprising 80% transcripts from reference genome annotation.

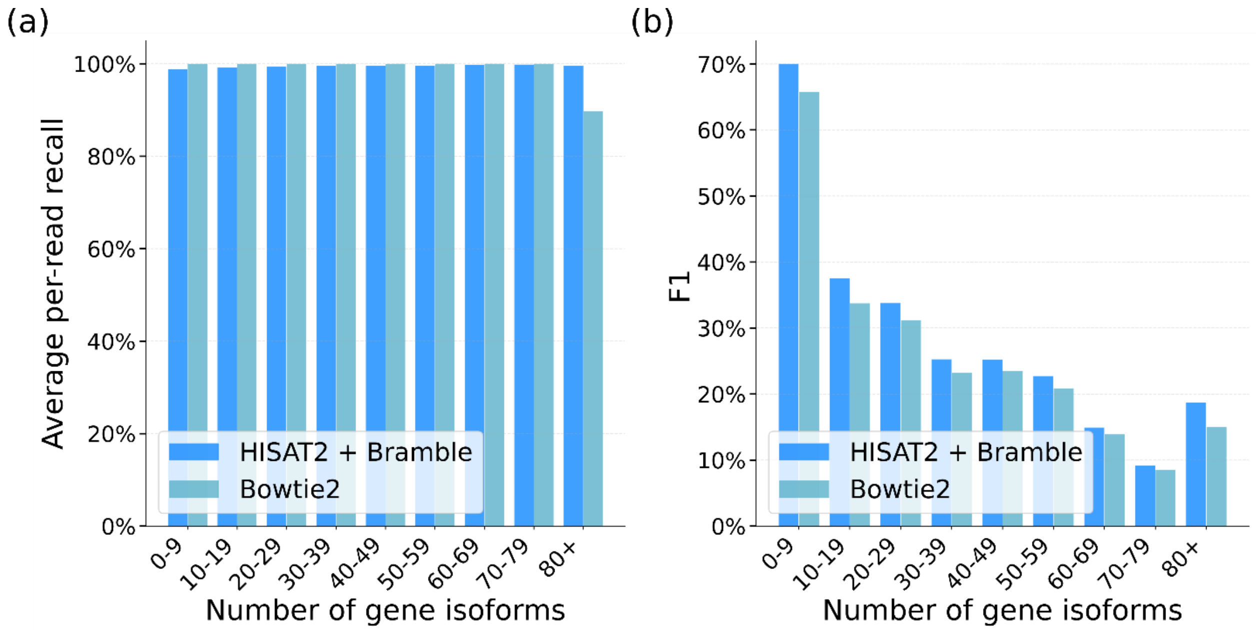


**Supplementary Figure 1.** Histograms showing a) average per-read recall and b) average F1 score per bin on simulated short-read data, binned by the number of gene isoforms belong to the gene of origin of the read. Per-read recall refers to a score of 1 if there is any alignment to the correct transcript of origin and 0 otherwise; scores are averaged across reads to get average per-read recall. F1 scores are calculated using average per-read precision and average per-read recall.

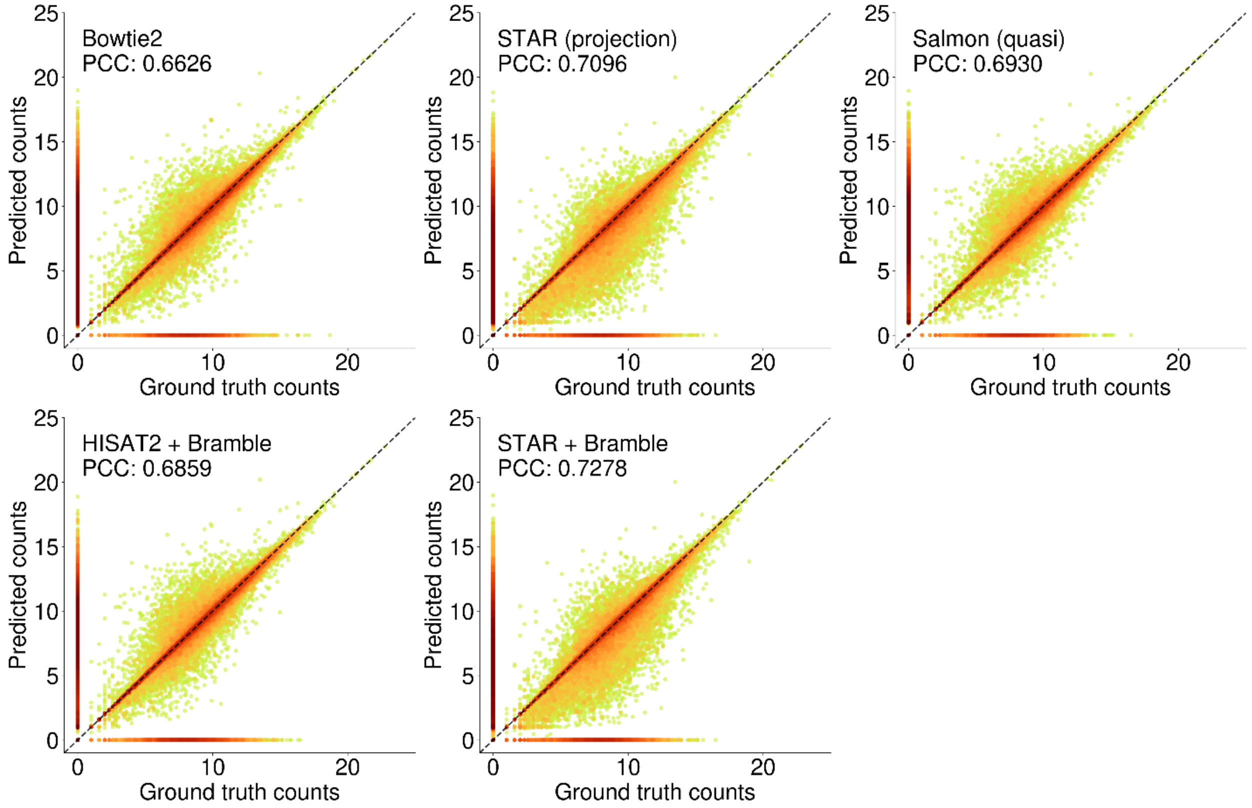


**Supplementary Figure 2.** Scatter plots showing log-scale ground truth counts vs. predicted counts on simulated short-read data for each method and the corresponding PCC values, using ground truth and corresponding abundance estimates from tissue 0, sample 0 for visualization. PCC is averaged across three tissues and 10 samples per tissue.

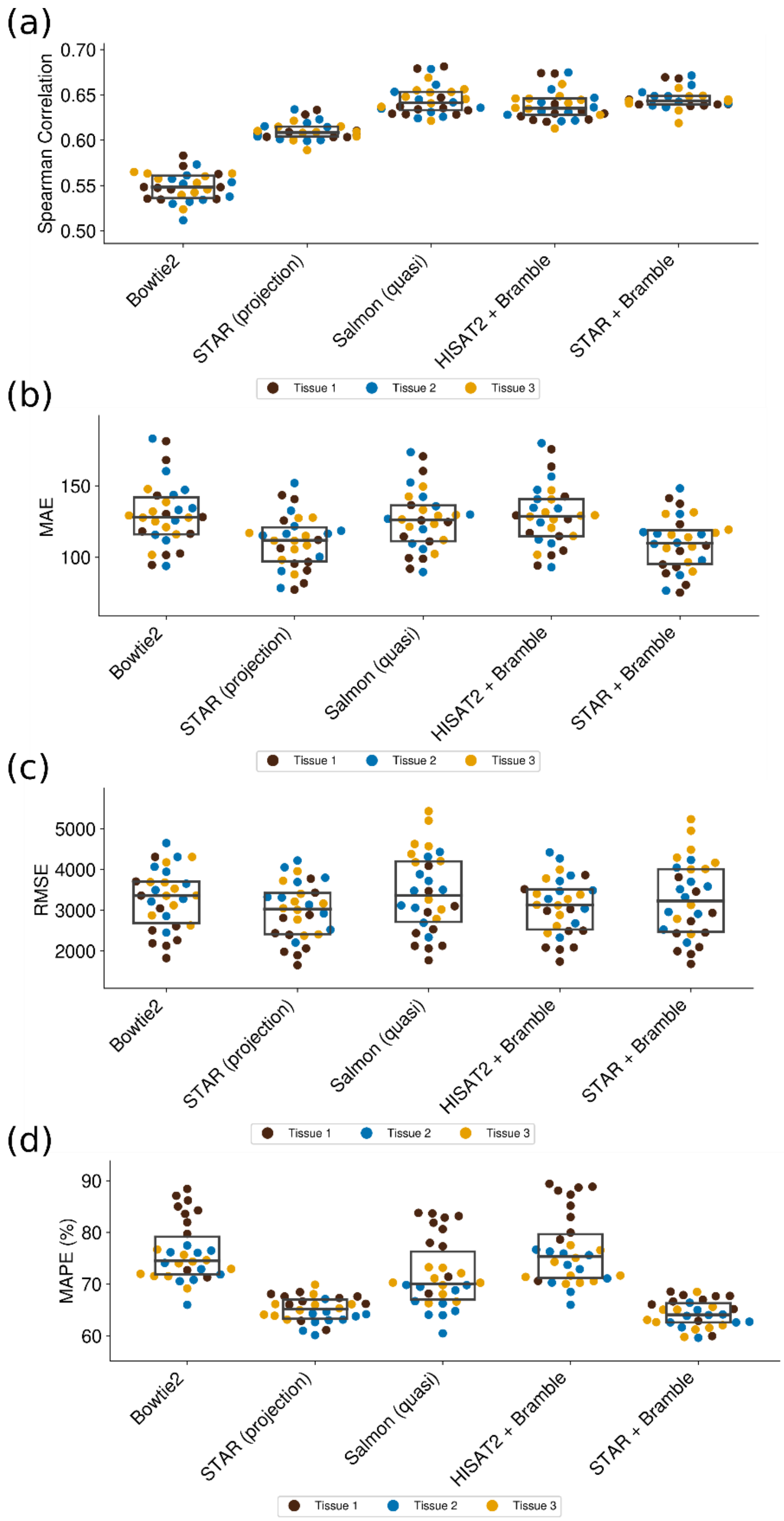


**Supplementary Figure 3.** Swarm plots showing a) Spearman Correlation (SCC), b) MAE, c) RMSE, d) MAPE on simulated short-read data. For boxes within swarms, interior lines indicate the median and the bottom and top ends indicate the first and third interquartile range. Metrics are averaged across three tissues and 10 samples per tissue.

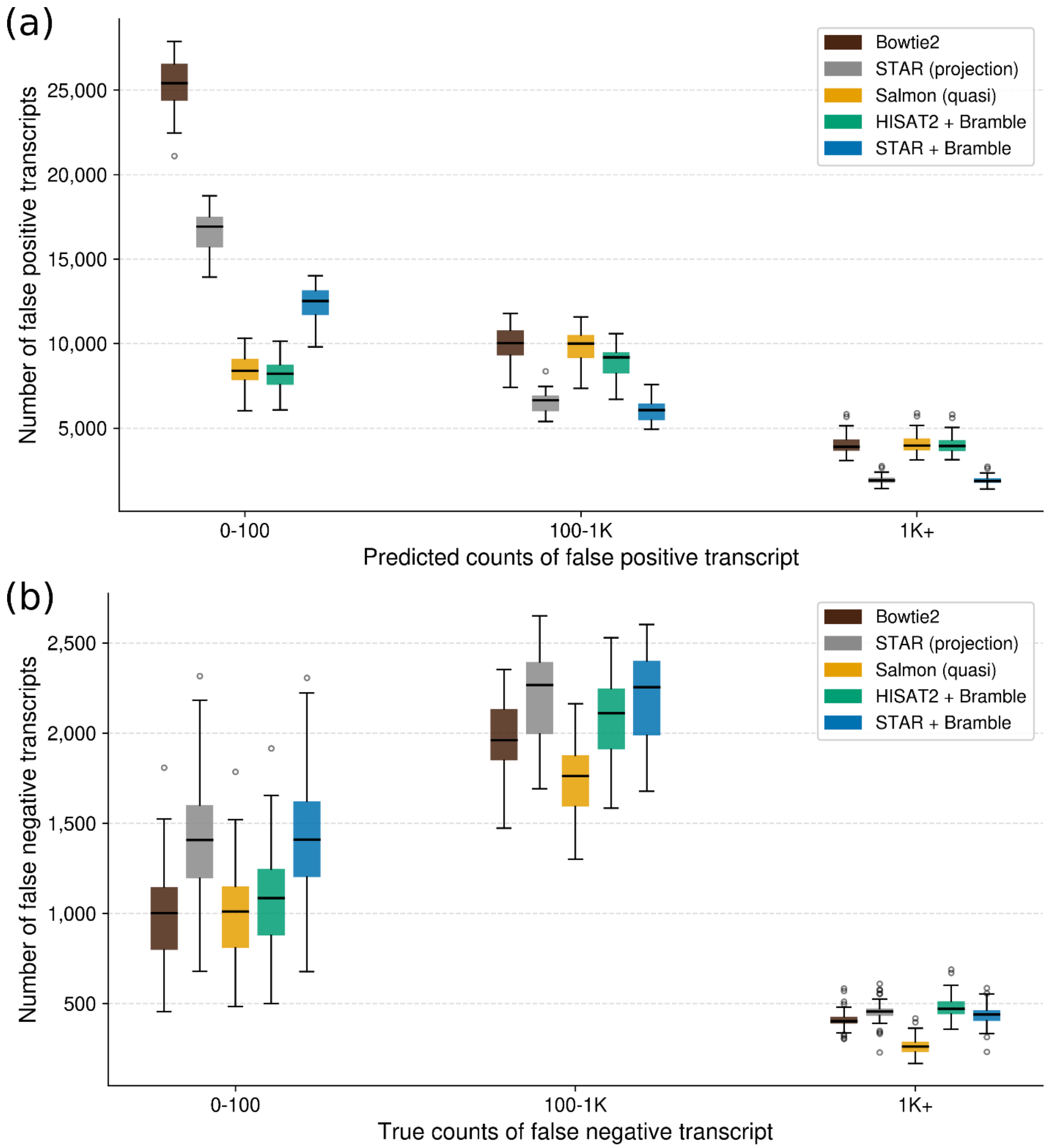


**Supplementary Figure 4.** Histograms showing a) false positives and b) false negatives on simulated short-read data. Values are averaged across three tissues and 10 samples per tissue.

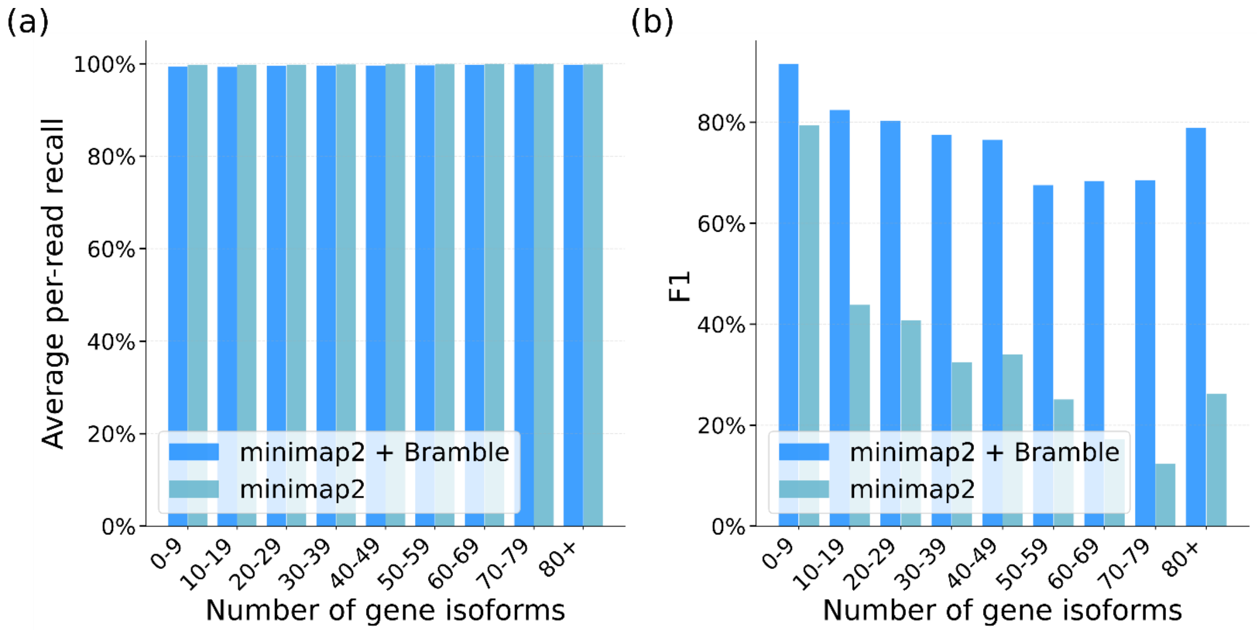


**Supplementary Figure 5.** Histograms showing a) average per-read recall and b) average F1 score per bin on simulated PacBio data, binned by the number of gene isoforms belong to the gene of origin of the read. Per-read recall refers to a score of 1 if there is any alignment to the correct transcript of origin and 0 otherwise; scores are averaged across reads to get average per-read recall. F1 scores are calculated using average per-read precision and average per-read recall.

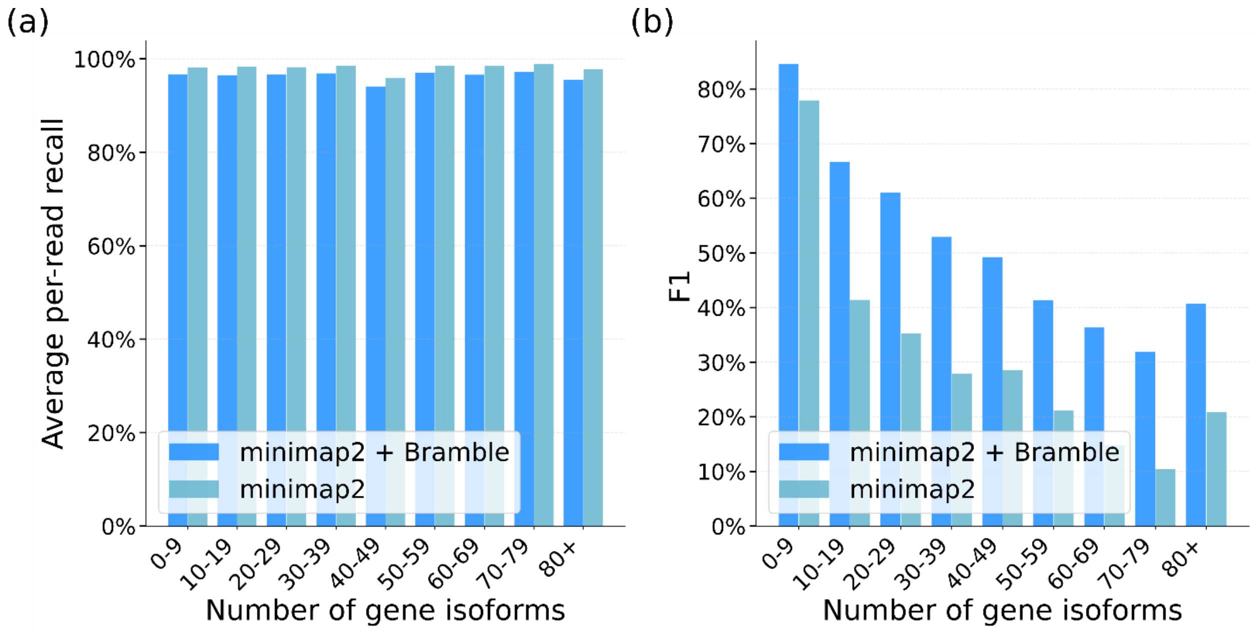


**Supplementary Figure 6.** Histograms showing a) average per-read recall and b) average F1 score per bin on simulated ONT data, binned by the number of gene isoforms belong to the gene of origin of the read. Per-read recall refers to a score of 1 if there is any alignment to the correct transcript of origin and 0 otherwise; scores are averaged across reads to get average per-read recall. F1 scores are calculated using average per-read precision and average per-read recall.

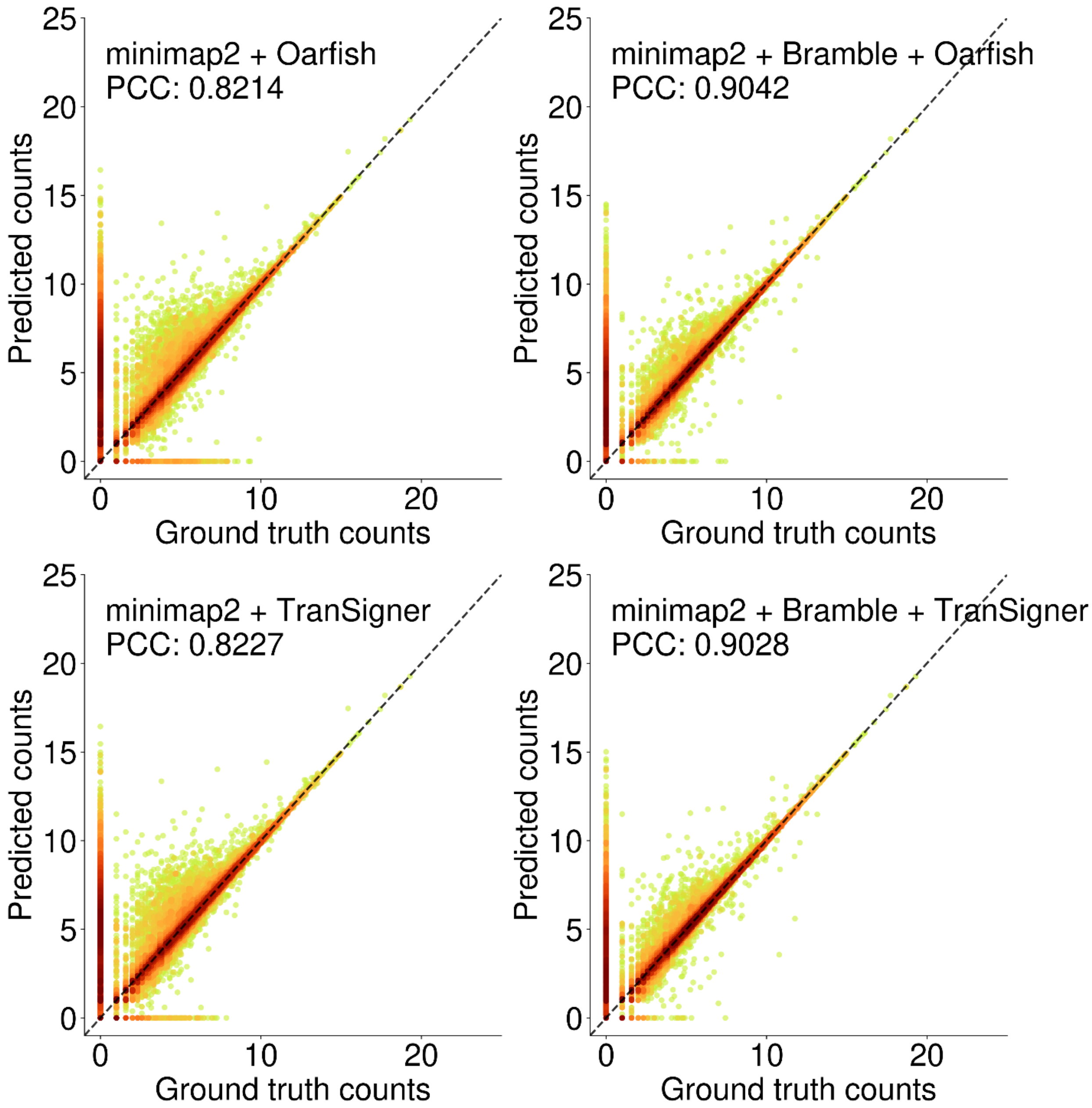


**Supplementary Figure 7.** Scatter plots showing log-scale ground truth counts vs. predicted counts on simulated PacBio data for each method and the corresponding PCC values, using ground truth and corresponding abundance estimates from tissue 0, sample 0 for visualization. PCC is averaged across three tissues and 10 samples per tissue.

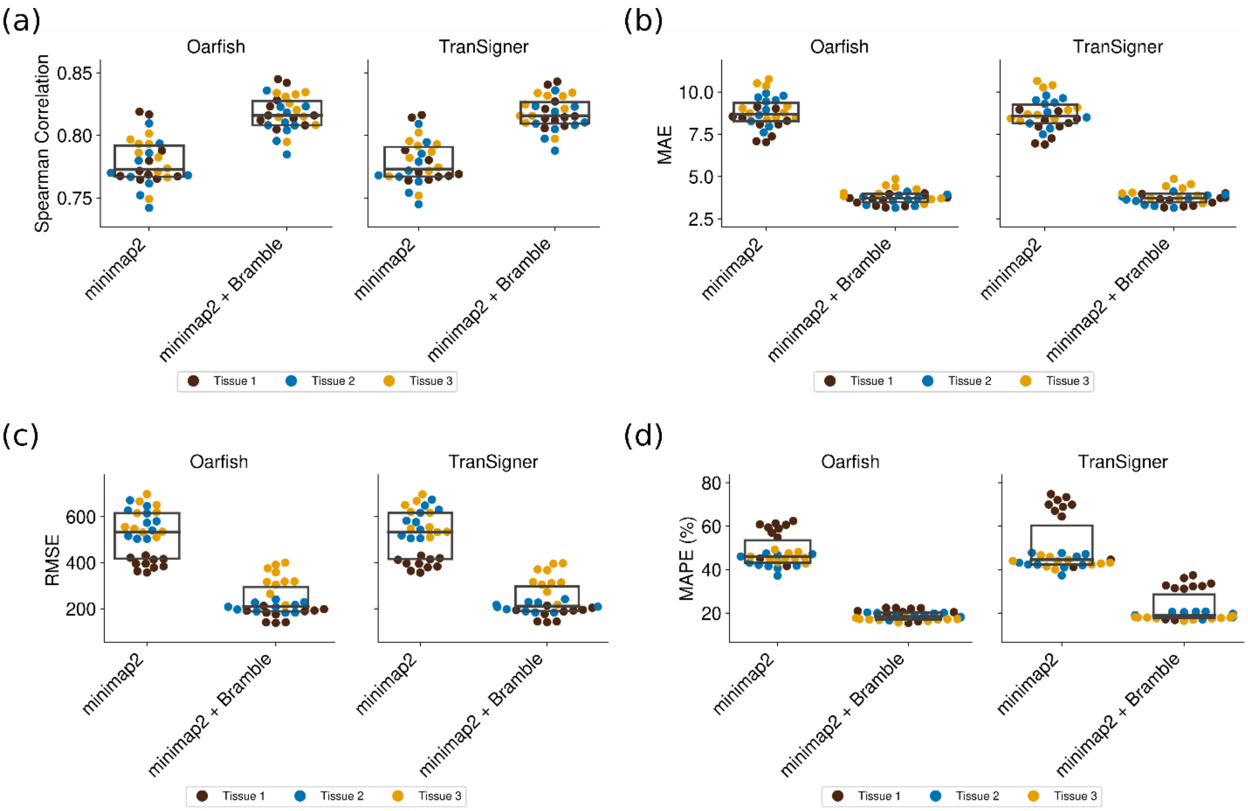


**Supplementary Figure 8.** Swarm plots showing a) Spearman Correlation (SCC), b) MAE, c) RMSE, d) MAPE on simulated PacBio data. For boxes within swarms, interior lines indicate the median and the bottom and top ends indicate the first and third interquartile range. Metrics are averaged across three tissues and 10 samples per tissue.

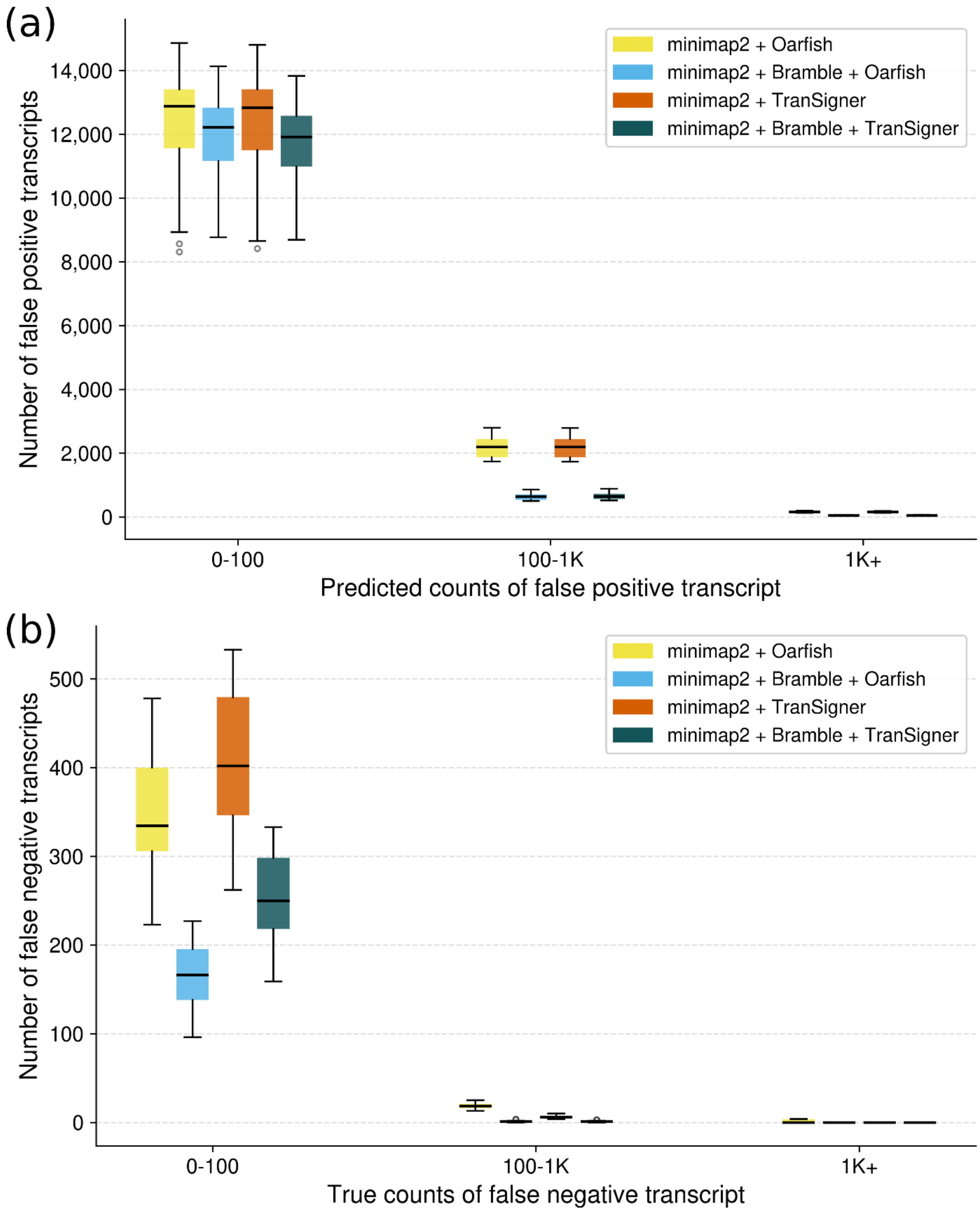


**Supplementary Figure 9.** Histograms showing a) false positives and b) false negatives on simulated PacBio data. Values are averaged across three tissues and 10 samples per tissue.

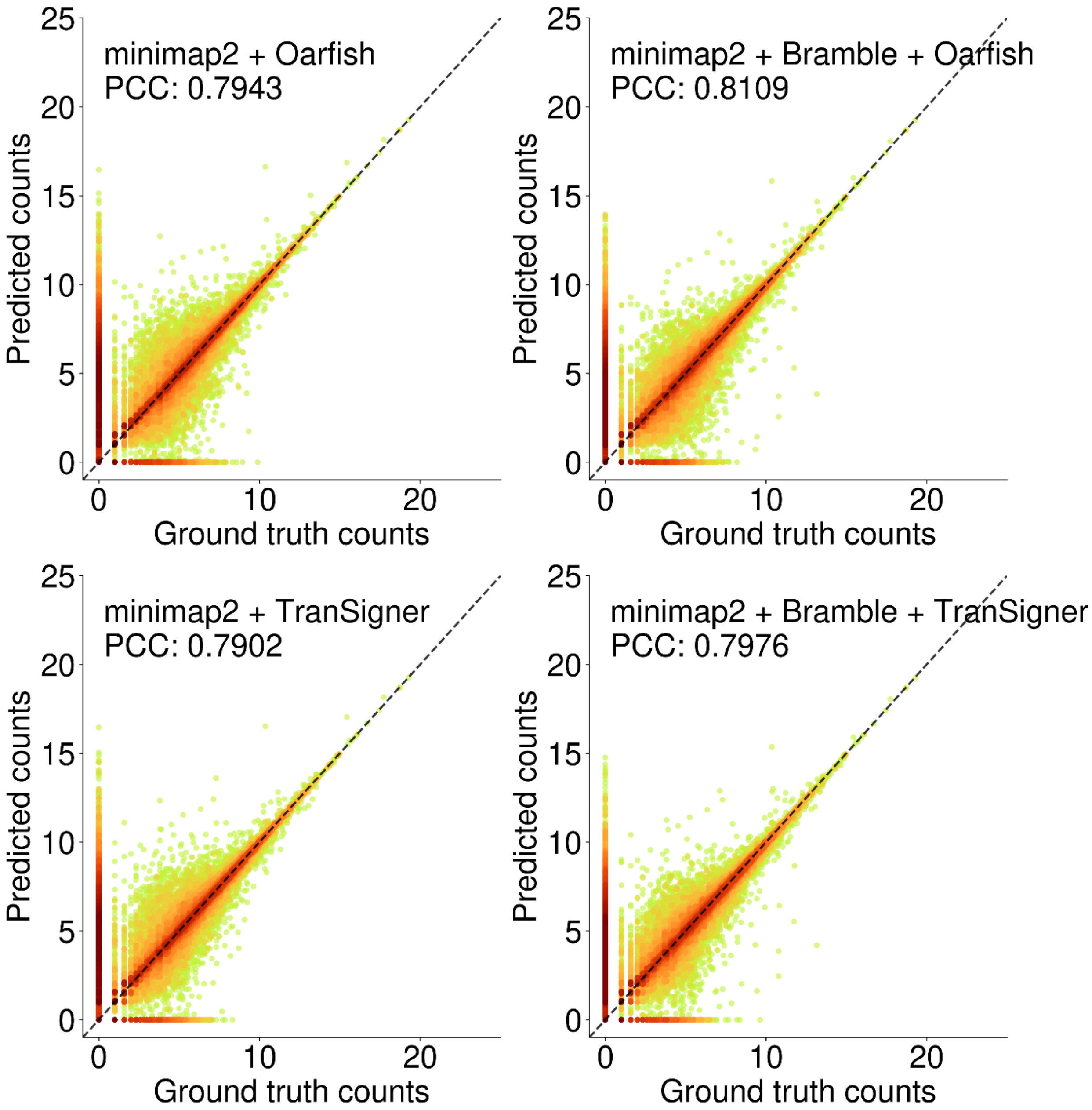


**Supplementary Figure 10.** Scatter plots showing log-scale ground truth counts vs. predicted counts on simulated ONT data for each method and the corresponding PCC values, using ground truth and corresponding abundance estimates from tissue 0, sample 0 for visualization. PCC is averaged across three tissues and 10 samples per tissue.

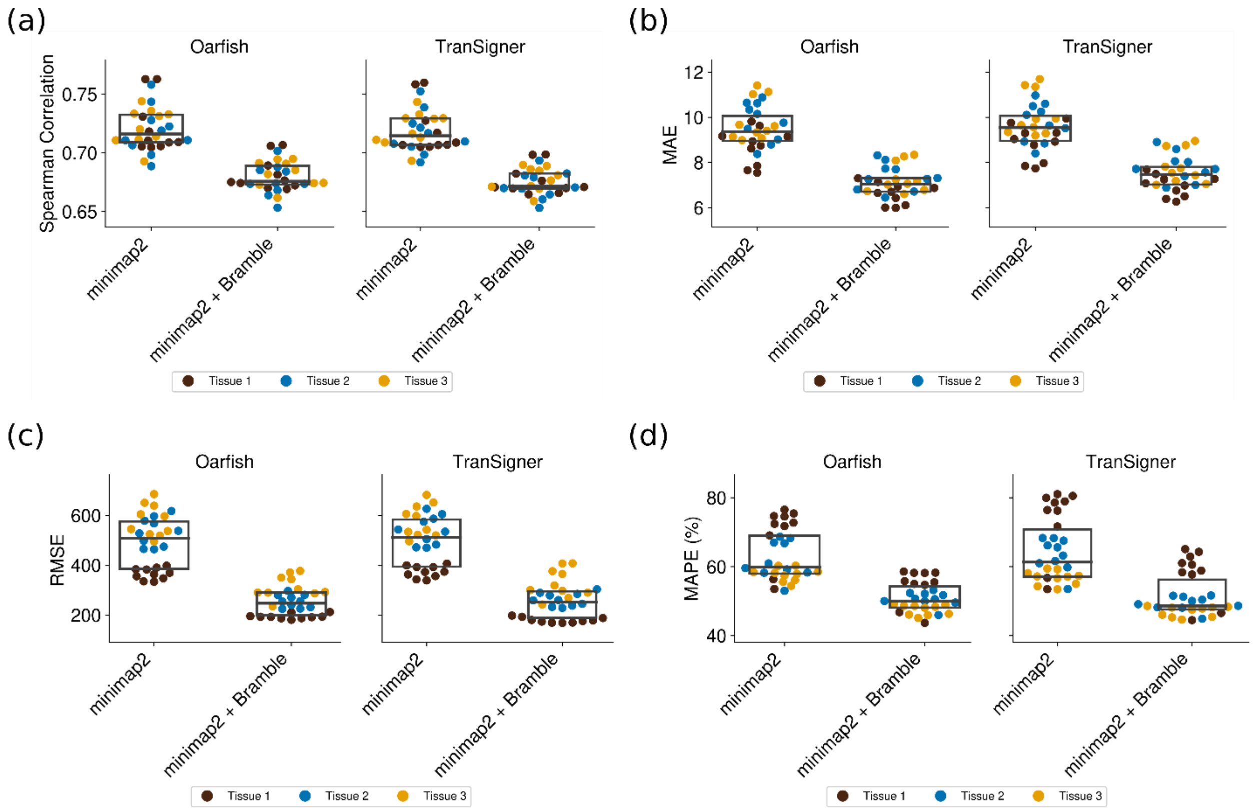


**Supplementary Figure 11.** Swarm plots showing a) Spearman Correlation (SCC), b) MAE, c) RMSE, d) MAPE on simulated ONT data. For boxes within swarms, interior lines indicate the median and the bottom and top ends indicate the first and third interquartile range. Metrics are averaged across three tissues and 10 samples per tissue.

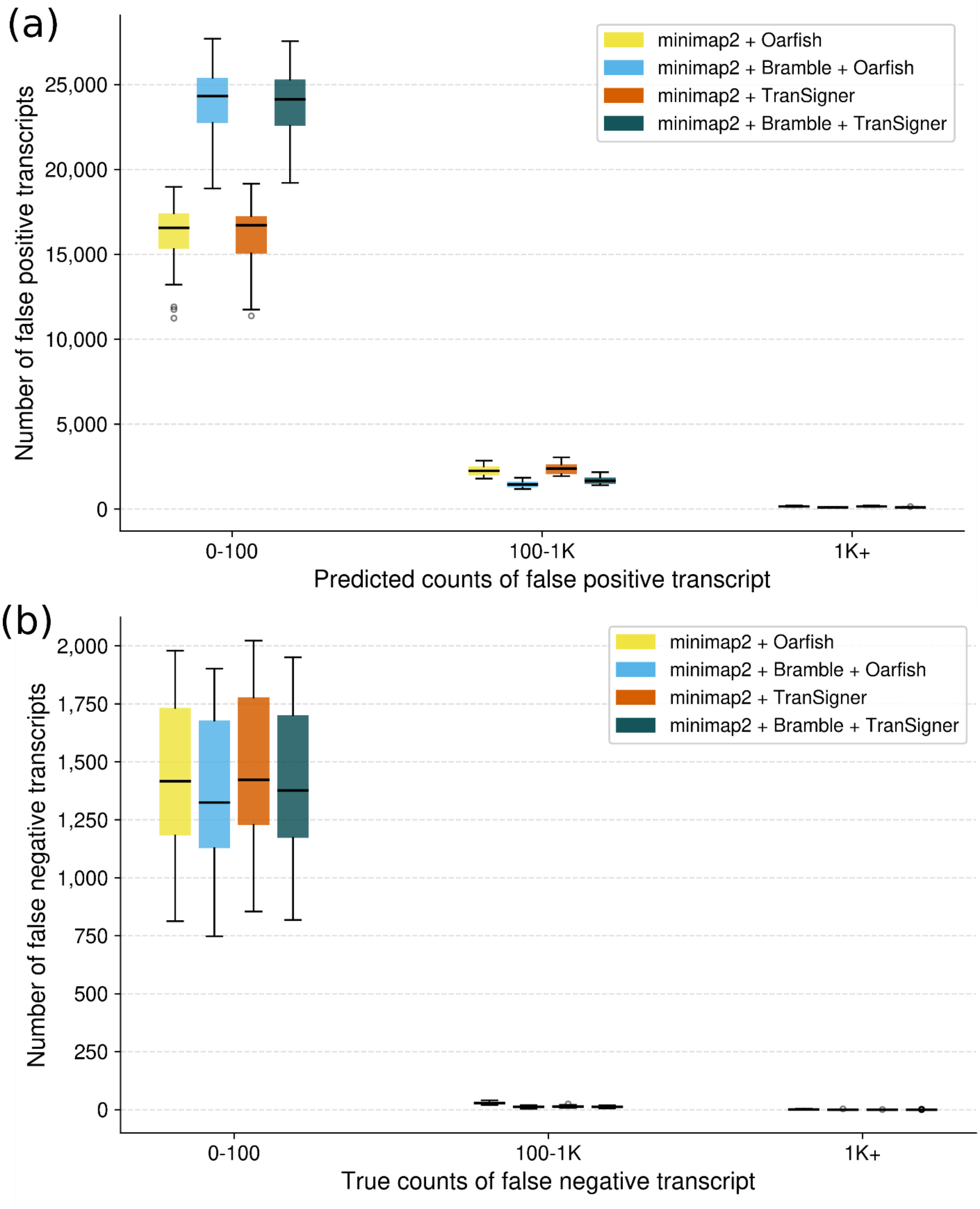


**Supplementary Figure 12.** Histograms showing a) false positives and b) false negatives on simulated ONT data. Values are averaged across three tissues and 10 samples per tissue.

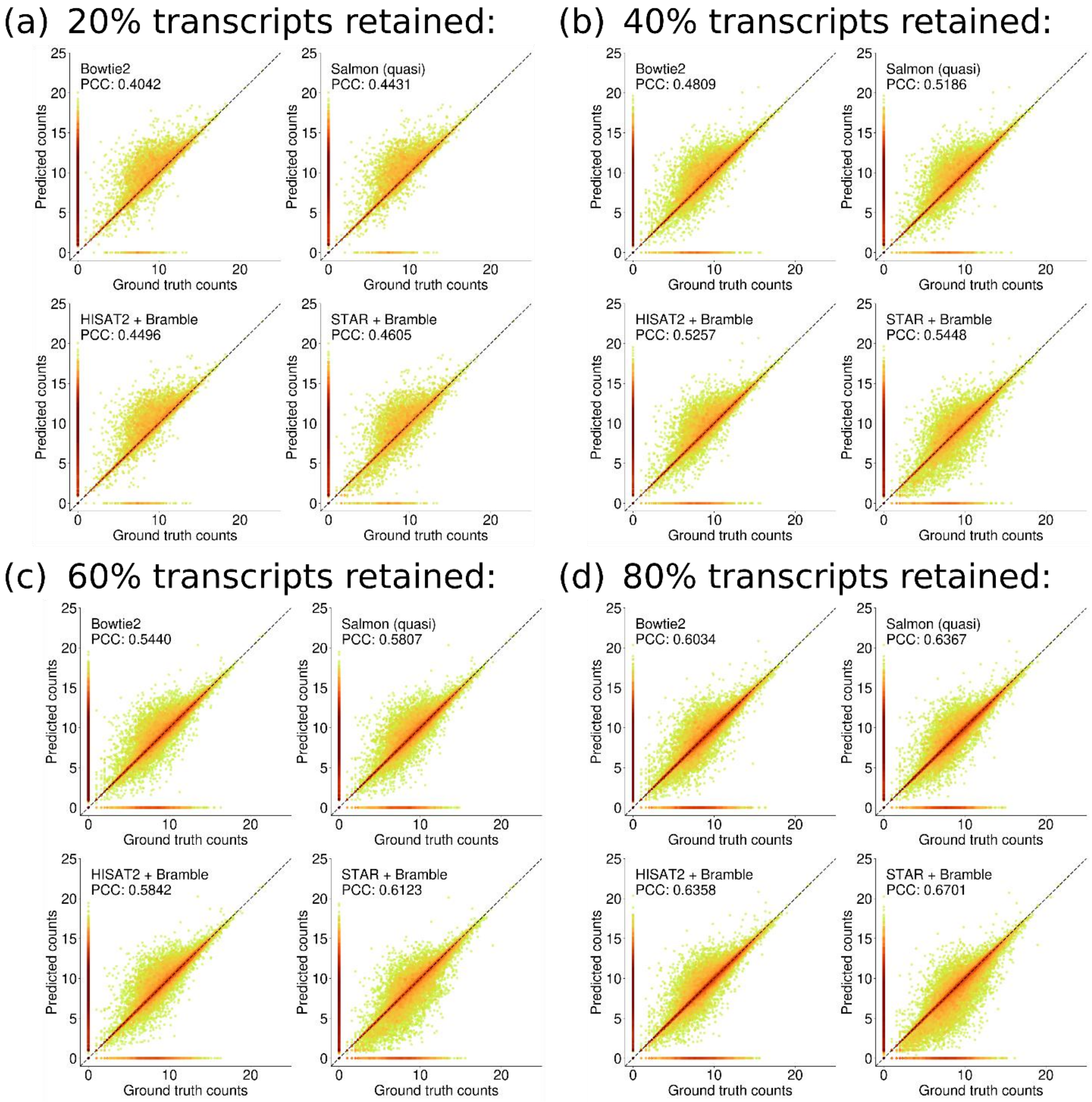


**Supplementary Figure 13.** Scatter plots showing log-scale ground truth counts vs. predicted counts on simulated short-read data with incomplete reference annotations for each method and the corresponding PCC values, using ground truth and corresponding abundance estimates from tissue 0, sample 0 for visualization. PCC is averaged across 10 samples from one simulated tissue. Incomplete reference annotations comprise a) 20%, b) 40%, c) 60%, and d) 80% transcripts from the human reference annotation.

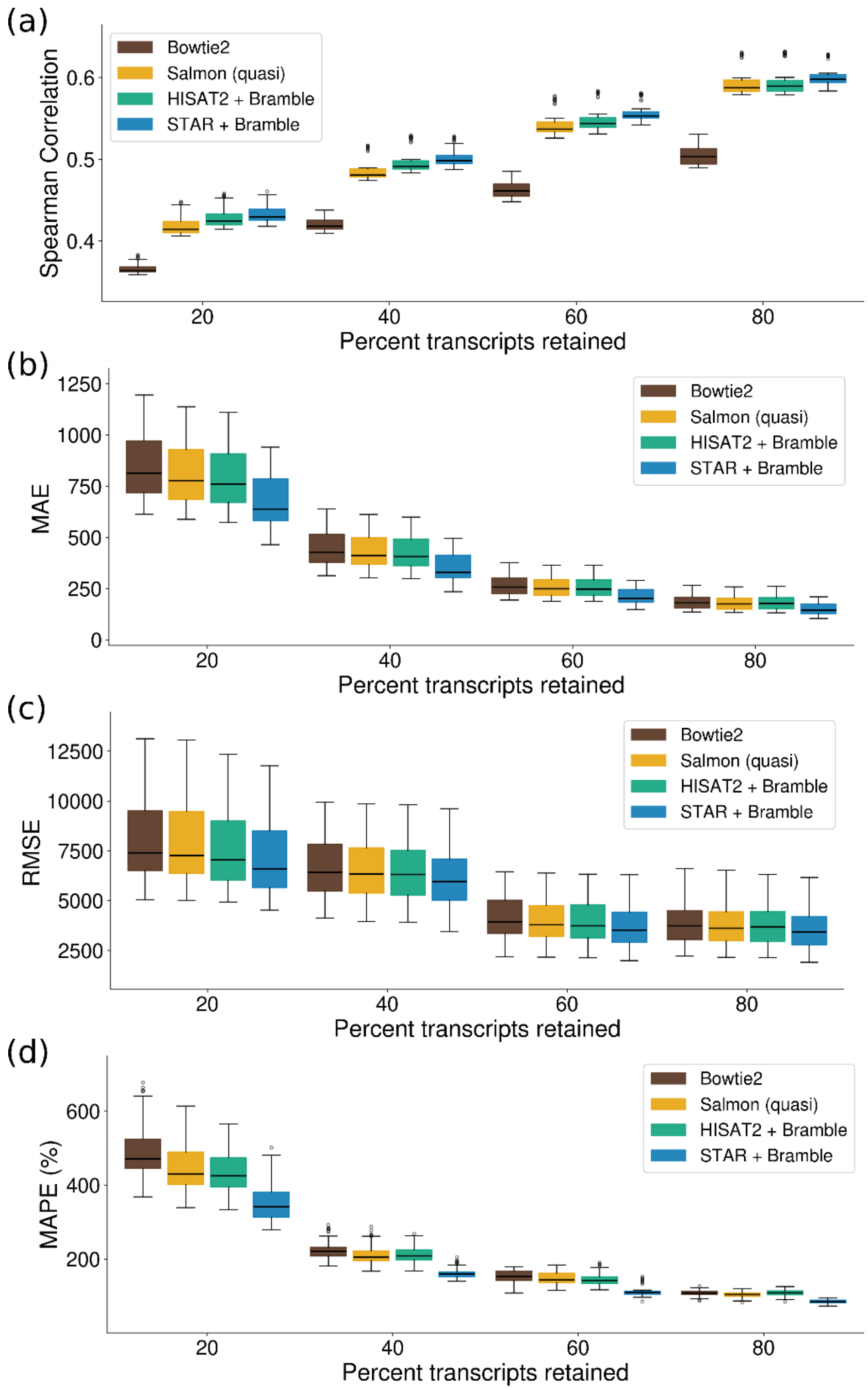


**Supplementary Figure 14.** Box-and-whisker plots showing a) Spearman Correlation (SCC), b) MAE, c) RMSE, d) MAPE on simulated short-read data with incomplete reference annotations. For boxes within swarms, interior lines indicate the median and the bottom and top ends indicate the first and third interquartile range. Metrics are averaged across 10 samples from one simulated tissue. Incomplete reference annotations comprise 20%, 40%, 60%, and 80% transcripts from the human reference annotation.

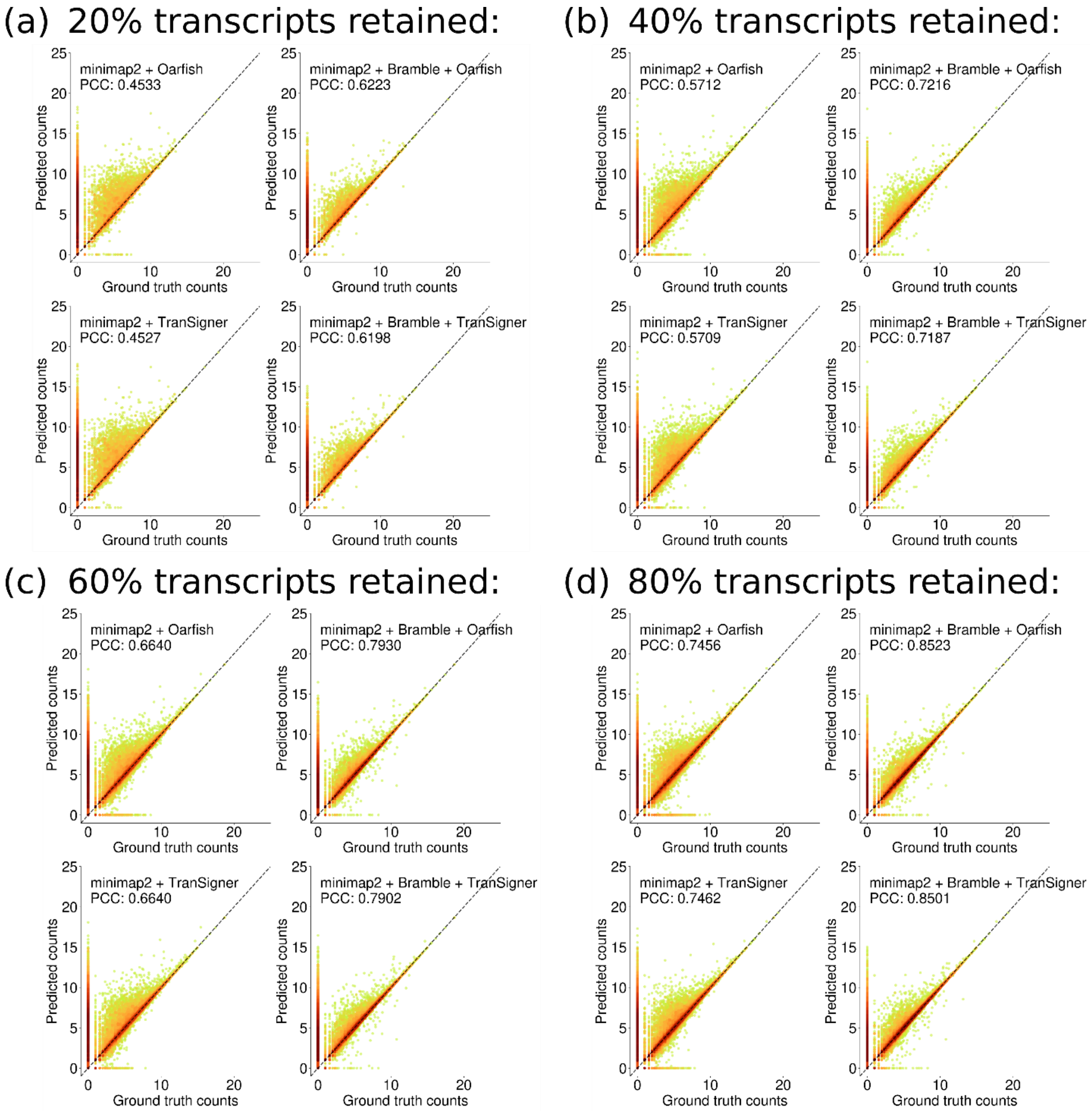


**Supplementary Figure 15.** Scatter plots showing log-scale ground truth counts vs. predicted counts on simulated PacBio data with incomplete reference annotations for each method and the corresponding PCC values, using ground truth and corresponding abundance estimates from tissue 0, sample 0 for visualization. PCC is averaged across 10 samples from one simulated tissue. Incomplete reference annotations comprise a) 20%, b) 40%, c) 60%, and d) 80% transcripts from the human reference annotation.

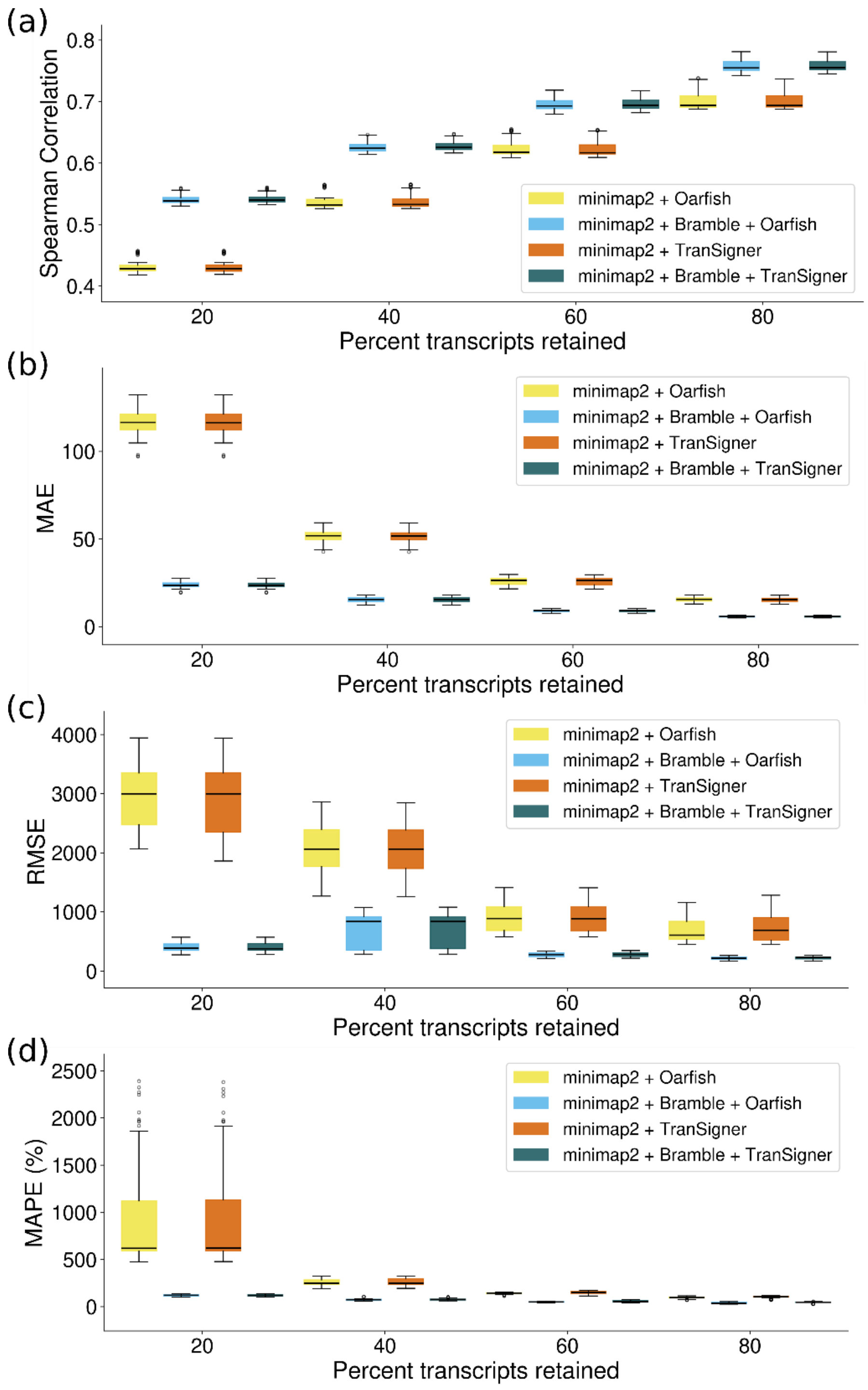


**Supplementary Figure 16.** Box-and-whisker plots showing a) Spearman Correlation (SCC), b) MAE, c) RMSE, d) MAPE on simulated PacBio data with incomplete reference annotations. For boxes within swarms, interior lines indicate the median and the bottom and top ends indicate the first and third interquartile range. Metrics are averaged across 10 samples from one simulated tissue. Incomplete reference annotations comprise 20%, 40%, 60%, and 80% transcripts from the human reference annotation.

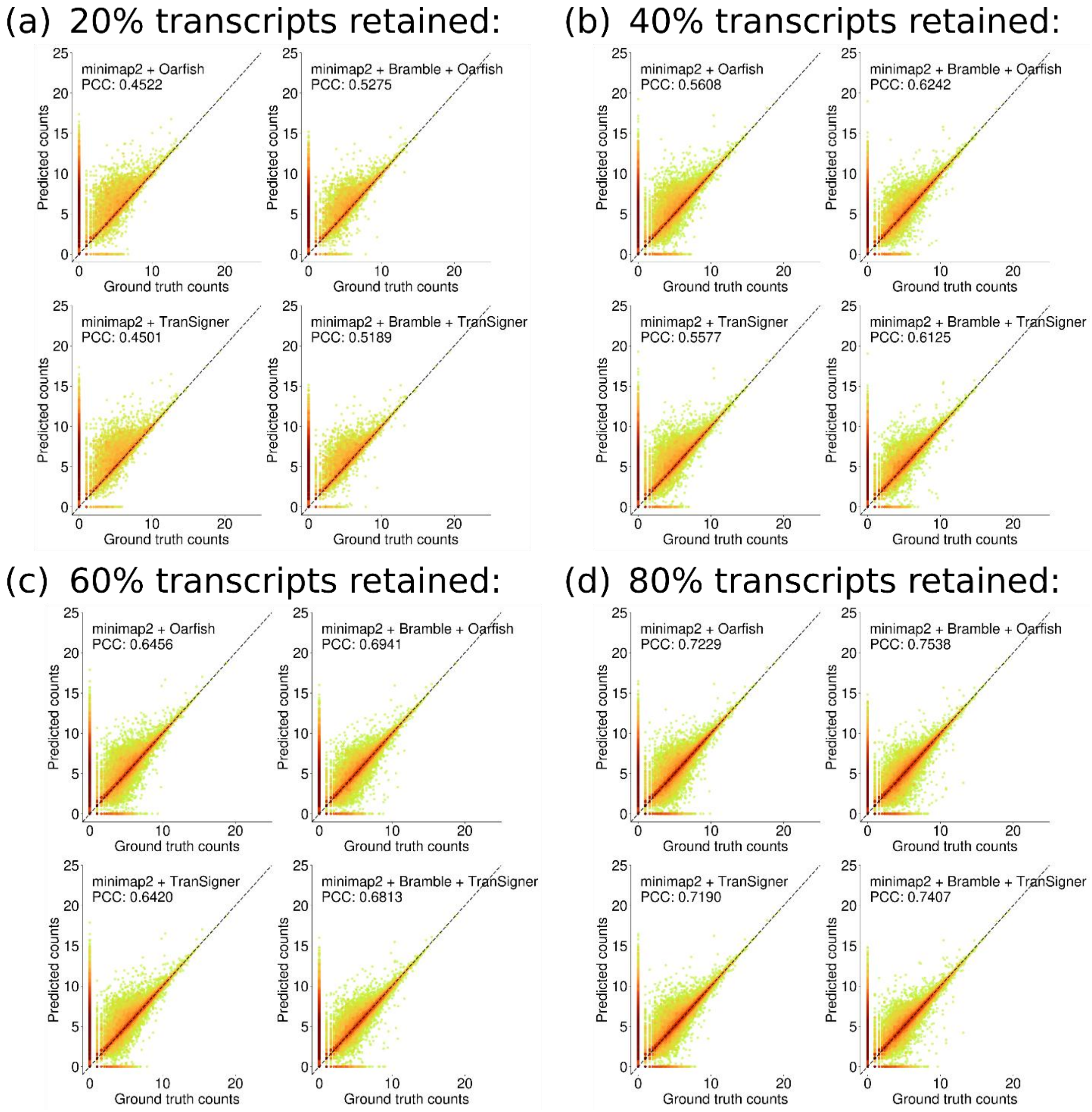


**Supplementary Figure 17.** Scatter plots showing log-scale ground truth counts vs. predicted counts on simulated ONT data with incomplete reference annotations for each method and the corresponding PCC values, using ground truth and corresponding abundance estimates from tissue 0, sample 0 for visualization. PCC is averaged across 10 samples from one simulated tissue. Incomplete reference annotations comprise a) 20%, b) 40%, c) 60%, and d) 80% transcripts from the human reference annotation.

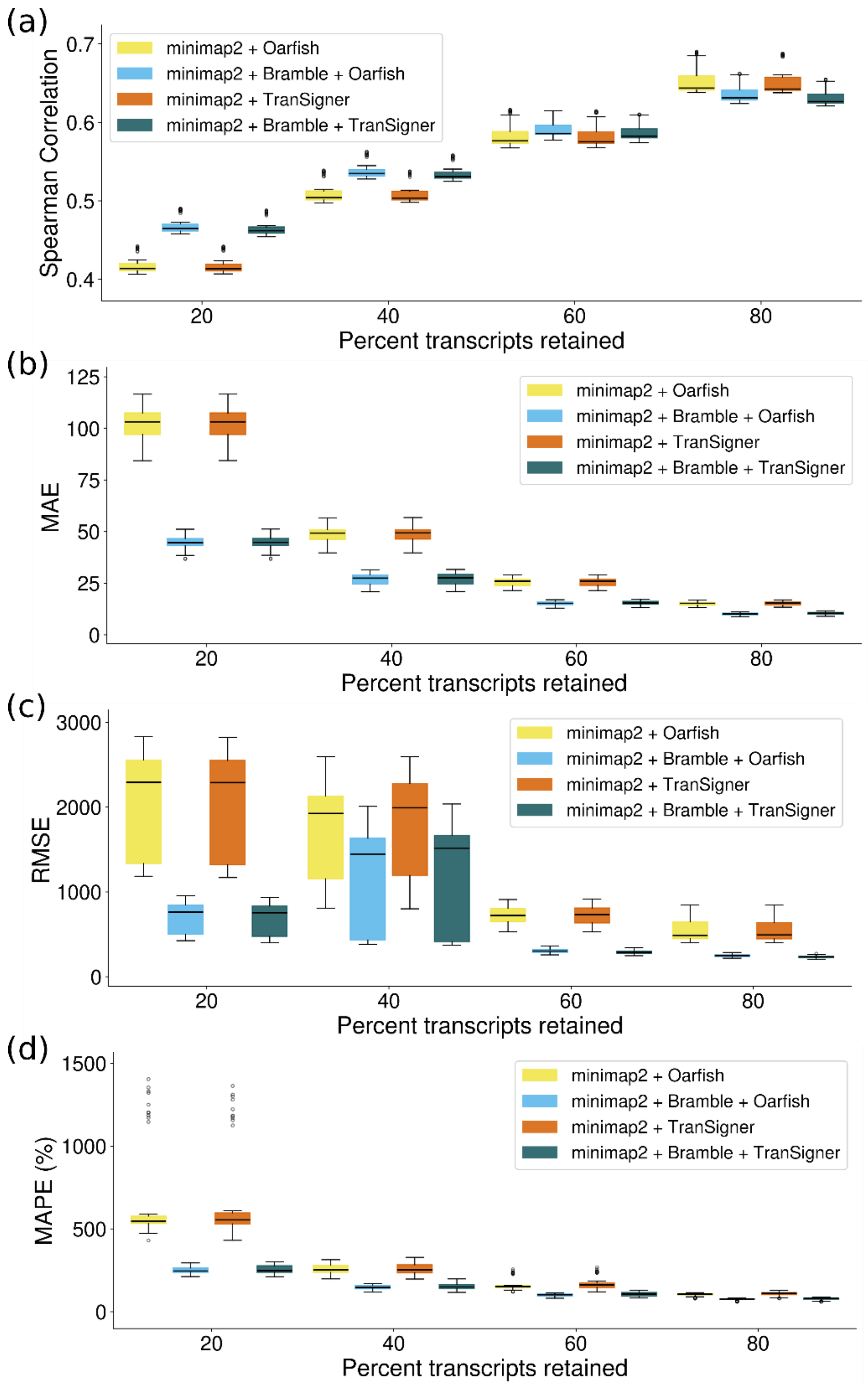


**Supplementary Figure 18.** Box-and-whisker plots showing a) Spearman Correlation (SCC), b) MAE, c) RMSE, d) MAPE on simulated ONT data with incomplete reference annotations. For boxes within swarms, interior lines indicate the median and the bottom and top ends indicate the first and third interquartile range. Metrics are averaged across 10 samples from one simulated tissue. Incomplete reference annotations comprise 20%, 40%, 60%, and 80% transcripts from the human reference annotation.

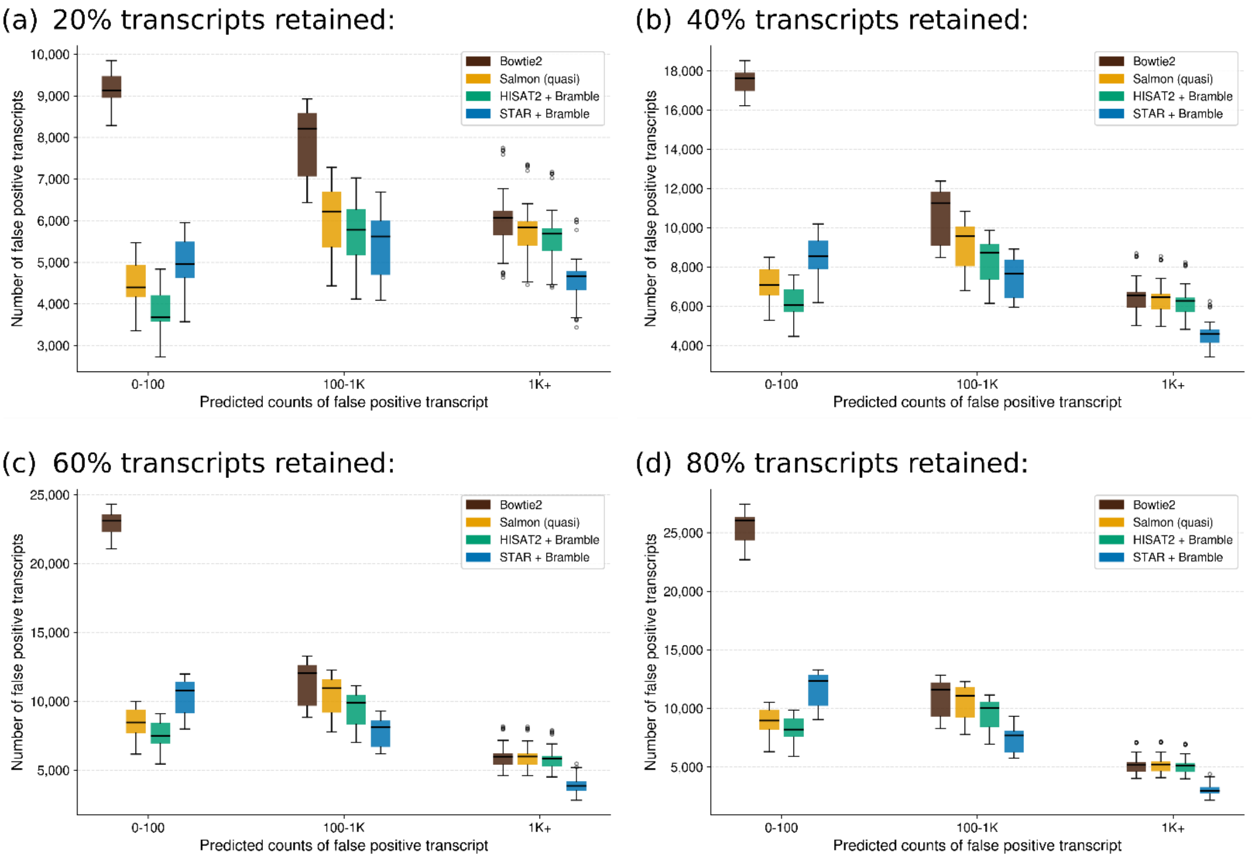


**Supplementary Figure 19.** Histograms showing false positives on simulated short-read data with incomplete reference annotations. Values are averaged across 10 samples from one simulated tissue. Incomplete reference annotations comprise a) 20%, b) 40%, c) 60%, and d) 80% transcripts from the human reference annotation.

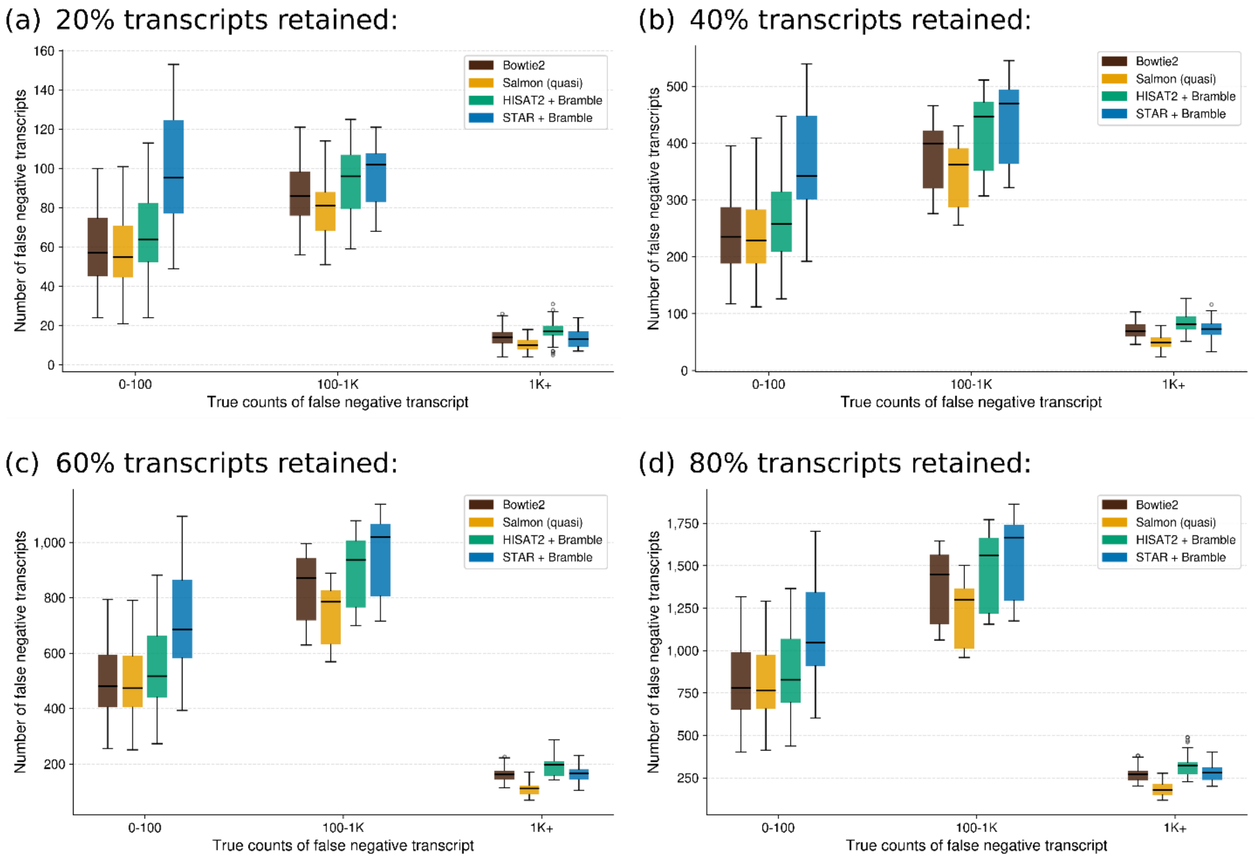


**Supplementary Figure 20.** Histograms showing false negatives on simulated short-read data with incomplete reference annotations. Values are averaged across 10 samples from one simulated tissue. Incomplete reference annotations comprise a) 20%, b) 40%, c) 60%, and d) 80% transcripts from the human reference annotation.

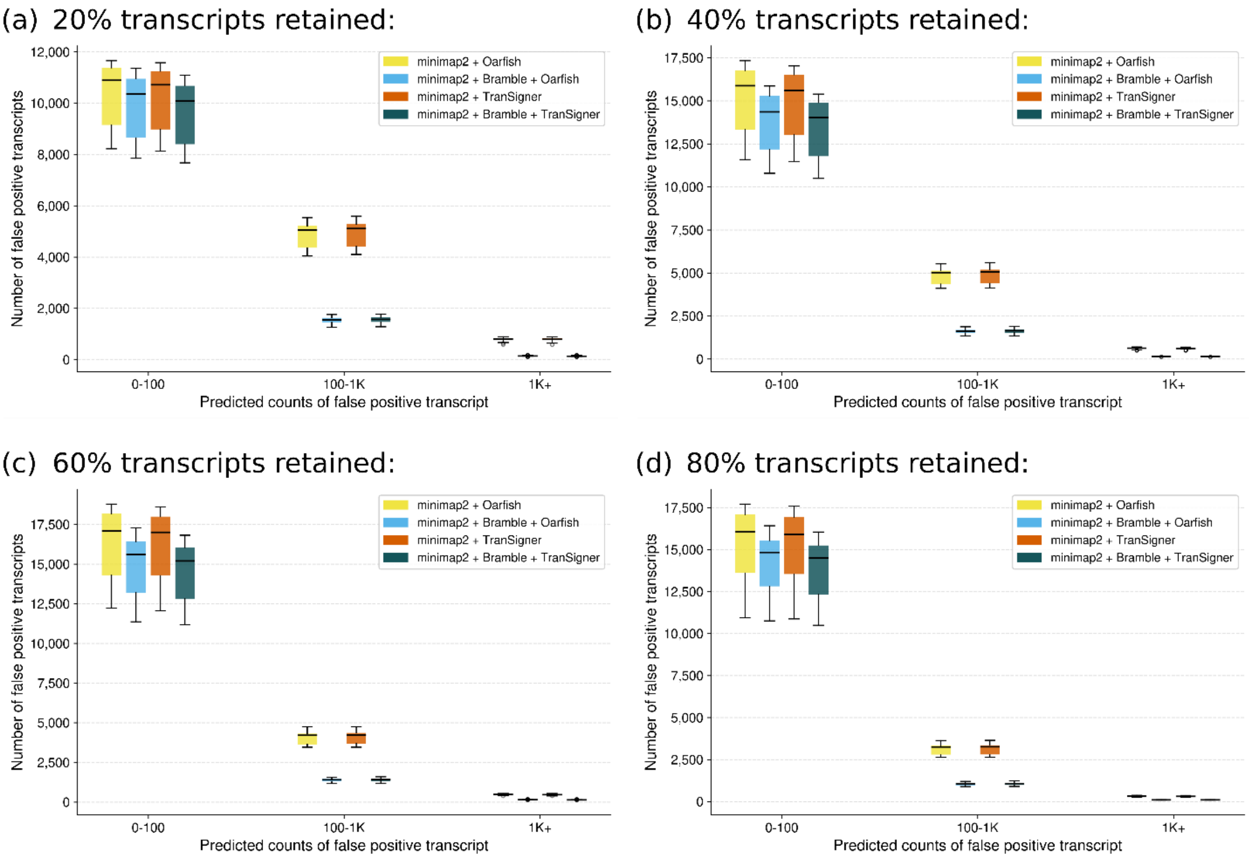


**Supplementary Figure 21.** Histograms showing false positives on simulated PacBio data with incomplete reference annotations. Values are averaged across 10 samples from one simulated tissue. Incomplete reference annotations comprise a) 20%, b) 40%, c) 60%, and d) 80% transcripts from the human reference annotation.

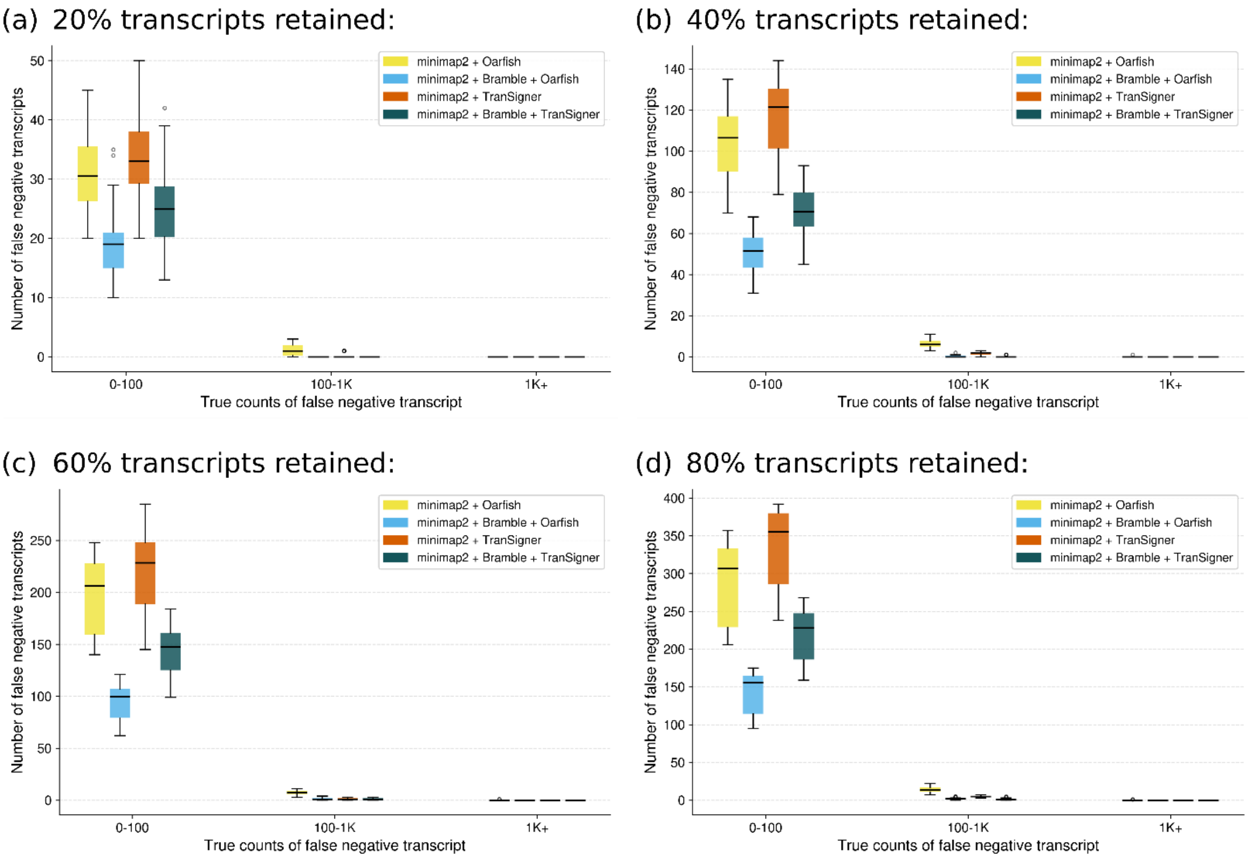


**Supplementary Figure 22.** Histograms showing false negatives on simulated PacBio data with incomplete reference annotations. Values are averaged across 10 samples from one simulated tissue. Incomplete reference annotations comprise a) 20%, b) 40%, c) 60%, and d) 80% transcripts from the human reference annotation.

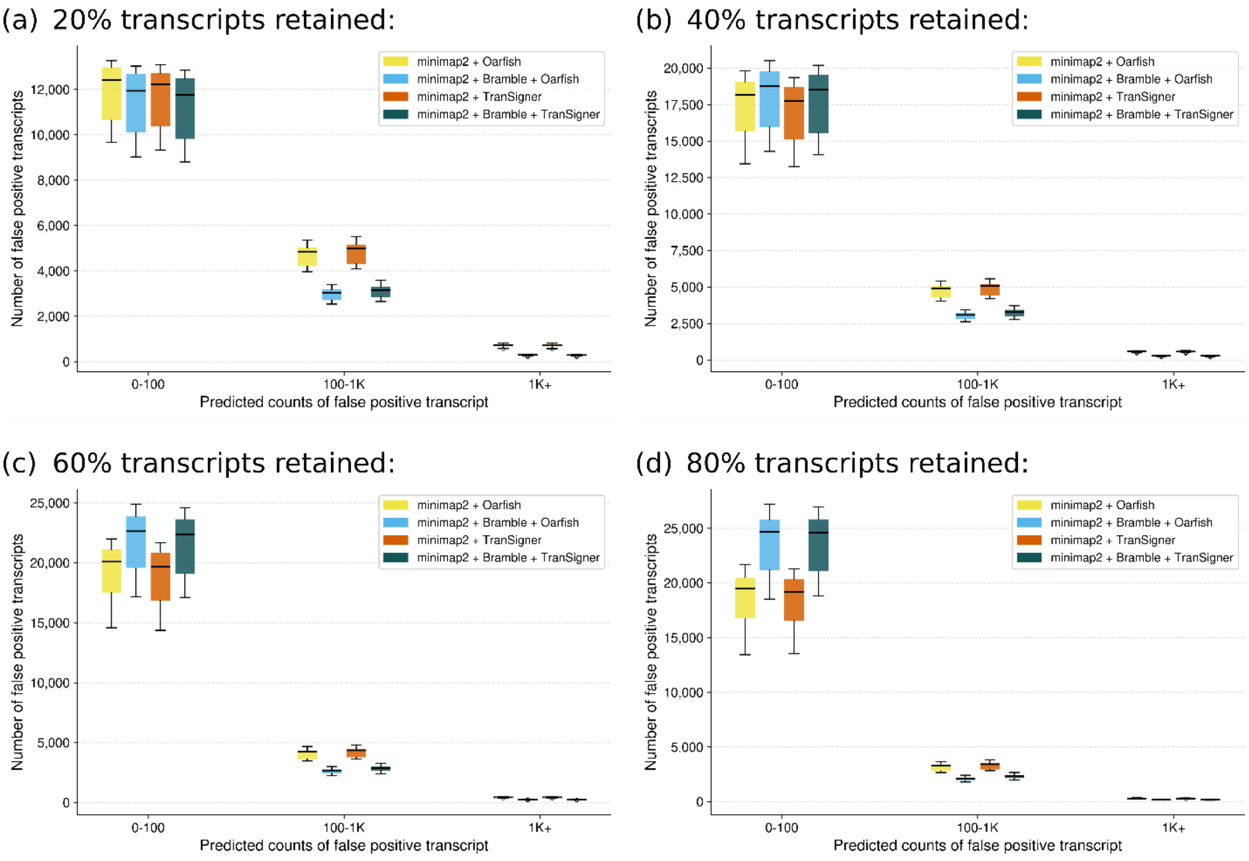


**Supplementary Figure 23.** Histograms showing false positives on simulated ONT data with incomplete reference annotations. Values are averaged across 10 samples from one simulated tissue. Incomplete reference annotations comprise a) 20%, b) 40%, c) 60%, and d) 80% transcripts from the human reference annotation.

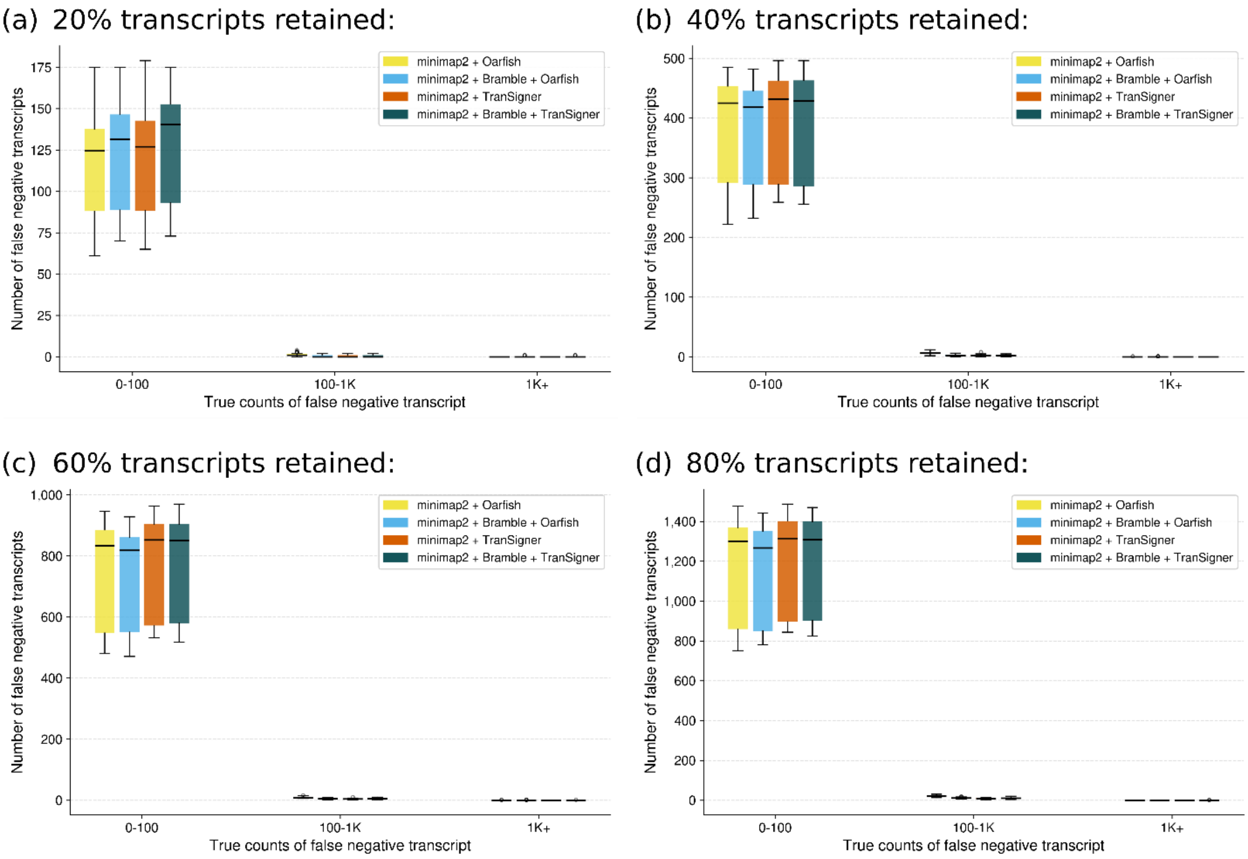


**Supplementary Figure 24.** Histograms showing false negatives on simulated ONT data with incomplete reference annotations. Values are averaged across 10 samples from one simulated tissue. Incomplete reference annotations comprise a) 20% b) 40% c) 60% and d) 80% transcripts from the human reference annotation.

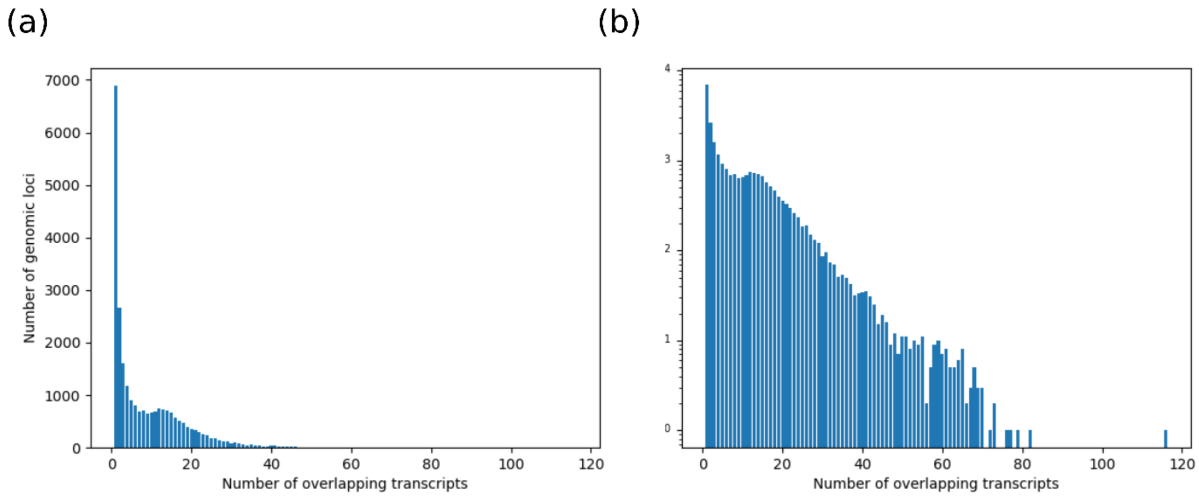


**Supplementary Figure 25.** Histograms showing the number of genomic loci with certain maximum numbers of overlapping transcripts, on a) a linear-scale and b) a log-scale.

**Supplementary Note 1. Command line usage of alignment and quantification tools**

For short-read alignment, we used Bowtie2 for transcriptome alignment, HISAT2 and STAR for genome alignment, and Salmon for quantification. We aligned reads to either the GRCh38 reference genome or the corresponding reference transcriptome, which we generated using `gffread` (Pertea and Pertea 2020) and the expanded CHESS 2.2 annotation. We built transcriptome indices for Bowtie2 and Salmon (building Salmon index decoys using the entire reference genome sequence), and genome indices for HISAT2 and STAR. To build indices, default parameters were used with the following exceptions:

- STAR index: `--sjdbGTFfile data/ref/chess2.2_ALL.gtf --sjdbOverhang 100`
- Salmon index: `-d decoys.txt`

To run transcriptome/genome alignment, default parameters were used with the following exceptions:

- Bowtie2: `-k 76`
- STAR: `--outSAMtype SAM --outStd SAM --sjdbOverhang 100 --quantMode TranscriptomeSAM`

We ran Bramble with default parameters on the genome-aligned BAM files produced by HISAT2 and STAR. To do transcript quantification, Salmon was run with `-l SF`.

For long-read alignment, we used minimap2 for transcriptome and genome alignment, and Oarfish and TranSigner for quantification. We used the same reference genome and transcriptome as described above. We created transcriptome and genome indices for minimap2 using default parameters. To align reads to the genome, minimap2 was run with:

- PacBio data: `-a -x splice:hq --junc-bed junc.bed`
- ONT data: `-a -x splice --junc-bed junc.bed`

To align reads to the transcriptome, minimap2 was run with:

- PacBio data: `-N 115 -a -x lr:hq`
- ONT data: `-N 115 -a -x map-ont`

We ran Bramble with `--lr -S GRCh38.fa` on the genome-aligned BAM files produced by minimap2. To do transcript quantification, we used the following settings:

- Oarfish: `--filter-group no-filters --model-coverage`
- TranSigner: `--keep-low --use-psw`

**Supplementary Note 2: Long-read clip rescue**

When junction-spanning reads are aligned to the genome and their terminal bases belong to the exons adjacent to the ones the alignments already reach, spliced genome aligners sometimes soft clip them instead of extending the alignments. This may occur when the anchor, the segment of the read extending into the adjacent exon, is too short to be confidently called as part of a spliced alignment. For long-reads, this can have detrimental effects on transcript quantification accuracy, because long-read quantification methods use alignment scores to rank candidate transcripts (see Supplementary Note 3). A soft-clipped exon is excluded from a candidate transcript's matched exon chain, lowering the alignment score and weakening its distinction from competing candidates. To address this, Bramble provides a clip-rescue mode available for long-read data. When enabled, clip rescue attempts to extend soft-clipped alignments into the neighboring ends of each candidate transcript, independently for the left and right ends of reads.

***Rescue criteria.*** Let $A$ be a genomic long-read alignment with left and right soft-clip lengths $n_L, n_R$, and let $t$ be a candidate transcript surviving the exon-chain matching procedure of Section 5.1.3, with terminal matched exons $e_1$ (leftmost) and $e_p$ (rightmost). Using the boundary offsets of Section 5.1.2, rescue is attempted only when:

- Side $L$: $n_L > 5$ and $\delta_L(a_1, e_1) \leq 0$ (i.e. the alignment already reaches $e_1$'s boundary without a deletion at the junction relative to the reference)
- Side $R$: $n_R > 5$ and $\delta_R(a_p, e_p) \leq 0$ (same as above)

***Query and reference sequence construction.*** The rescue query $q$ is the concatenation of the read's clipped terminus with any boundary insertion recorded at the matched exon:

- Side $L$: $|q| = n_L + max(-\delta_L(a_1, e_1), 0)$
- Side $R$: $|q| = n_R + max(-\delta_R(a_p, e_p), 0)$

A reference sequence $g$ is assembled by querying the genome-to-transcriptome tree for the exons of $t$ immediately preceding (left) or following (right) $e_1$ or $e_p$, concatenating their sequences outward until $|g| \geq |q|$. Rescue is abandoned if $t$ has no neighboring exon on that side.

***Extension alignment.*** A window of $g$ of length $|q| + 40$, taken from the end nearest to the matched terminal exon, is aligned against $q$ using ksw2's ksw_extz2_sse routine (Suzuki and Kasahara 2018; Li 2018) in extension mode (`KSW_EZ_EXTZ_ONLY`) with approximate maximum score tracking (`KSW_EZ_APPROX_MAX, KSW_EZ_APPROX_DROP`), an affine gap penalty (match +1, mismatch -4, gap open 4, gap extend 1), and a Z-drop threshold of 40. For left-side rescue, both $q$ and the reference window are reversed before alignment, so that ksw2's extension runs outward from the exon boundary in the read's original orientation. Let $s$ denote the resulting maximum extension score; rescue is abandoned if $s < 10$ or if the alignment fails ($s = -\infty$).

***Segment construction.*** On a successful rescue, any boundary insertion previously recorded at the matched exon is discarded, superseded by the rescue alignment. A new segment spanning the reference interval consumed by the alignment is prepended (left) or appended (right) to the exon chain matched between $A$ and $t$, retaining any unreached query bases as an ordinary soft clip, and is incorporated into $t$'s alignment CIGAR during CIGAR construction.

**Supplementary Note 3: Alignment score**

A revised alignment score is introduced to rank candidate alignments during quantification, extending the alignment score (AS) defined in the SAM specification. The alignment score is particularly important for long-read quantification, where it serves as a primary criterion for distinguishing among secondary alignments.

***Similarity score.*** Let $A$ be a genomic alignment evaluated in the context of a candidate transcript $t$. Bramble evaluates $A$ as a chain of exons matched to $t$; within each matched exon, $o(A)$ accumulates the interior matched bases, together with any bases inserted or deleted relative to $t$ at the exon's boundaries, analogous to the boundary offsets $\delta_L, \delta_R$ of Section 5.1.2, including any soft-clipped bases at the ends of $A$. Let $c(A, t)$ denote only the interior matched bases among these. The ratio

$$\rho(A, t) = \frac{c(A, t)}{o(A)}$$

reflects the structural consistency of $A$ with $t$: exon boundaries that diverge from $t$'s annotated splice sites inflate $o(A)$ without contributing to $c(A, t)$, lowering $\rho(A, t)$ accordingly. Let $J(A, t)$ denote the number of splice-junction boundaries in $A$ that coincide with splice sites described by $t$, plus one. The similarity score is then

$$S(A, t) = J(A, t) \cdot \left(\frac{\rho(A,t) - T}{1 - T}\right)^2.$$

Alignments spanning more junctions consistent with $t's$ annotated splice sites receive higher scores, reflecting greater confidence that $A$ represents a true isoform-level mapping rather than a spurious or ambiguous placement. If similarity filtering is enabled, candidates with $\rho(A, t) < T$, for a fixed similarity threshold $T$, are considered insufficiently supported and discarded; when similarity filtering is disabled, $S(A, t)$ is set to 1.

***Revised alignment score.*** The revised alignment score $AS'$ is

$$AS'(A, t) = S(A, t) \cdot (AS(A) + C(A)),$$

where $AS(A)$ is the alignment score as defined by the SAM specification, and $C(A)$ is the clip-rescue score, the cumulative score of successful alignments recovered by long-read clip rescue for $A$. When clip rescue is not used, $C(A) = 0$.

**Supplementary Note 4: Mapping quality**

Bramble calculates the mapping quality (MAPQ), a field defined by the SAM specification, using the same convention employed by aligners such as STAR (Dobin et al. 2012) and TopHat (Kim et al. 2013). For a read with $N_{map}$ recorded alignments, uniquely mapping reads ($N_{map} = 1$) are assigned $MAPQ = 255$, and multi-mapping reads are assigned

$$MAPQ = \left\lfloor -10 \cdot log_{10}(1 - \frac{1}{N_{map}}) \right\rfloor.$$

This ensures Bramble's output matches the MAPQ convention that downstream software expects.

**Supplementary Note 5: Paired-end reads**

Bramble applies a set of rules, in order of precedence, to determine how paired-end reads are handled at the transcript level. If the two mates map to different chromosomes, pairing information is discarded and alignments from each mate are processed independently. For pairs mapping to the same reference sequence:

1. If the mates' candidate transcript sets share one or more transcripts, only the shared transcripts are retained, since a valid transcript of origin must account for both mates.
2. If the mates share no transcripts, both are retained as a discordant mapping only when each mate maps to exactly one distinct transcript; otherwise, the pair is discarded, preventing exponential growth in reported alignments when the ambiguity cannot be resolved.
3. If either mate produces no match, the pair is discarded.

**References for supplementary materials**

1. Pertea, G. & Pertea, M. GFF Utilities: GffRead and GffCompare. *F1000Res* **9**, 10.12688/f1000research.23297.2. eCollection 2020 (2020).

2. Suzuki, H. & Kasahara, M. Introducing difference recurrence relations for faster semi-global alignment of long sequences. *BMC Bioinformatics* **19**, 45 (2018).

3. Li, H. Minimap2: pairwise alignment for nucleotide sequences. *Bioinformatics* **34**, 3094–3100 (2018).

4. Dobin, A. *et al*. STAR: ultrafast universal RNA-seq aligner. *Bioinformatics* **29**, 15–21 (2012).

5. Kim, D. *et al*. TopHat2: accurate alignment of transcriptomes in the presence of insertions, deletions and gene fusions. *Genome Biol* **14**, R36 (2013).
